# Paired Associates Learning is Disrupted After Unilateral Parietal Lobe Controlled Cortical Impact in Rats: A Trial-by-Trial Behavioral Analysis

**DOI:** 10.1101/2022.04.05.487213

**Authors:** Samantha M. Smith, Elena L. Garcia, Caroline Davidson, John Thompson, Sarah Lovett, Nedi Ferekides, Quinten Federico, Argyle V. Bumanglag, Abbi R. Hernandez, Jose F. Abisambra, Sara N. Burke

**Author notes:** Indicates corresponding author **Correspondence should be addressed to:** Sara N. Burke, PhD, Department of Neuroscience, University of Florida, P.O. Box 100244, Gainesville, FL 32611. **Author Contributions:** SMS collected data, contributed analytic tools, analyzed data, and wrote the manuscript. EG collected and analyzed data, CD collected and analyzed data, JT collected data, SL performed the surgeries, NF collected and analyzed data, QF collected data, AVB helped oversee and perform research, AH helped with experimental design and performed research, JFA and SNB designed the research and edited the manuscript.

## Abstract

Approximately 60-70 million people suffer from a traumatic brain injury (TBI) each year. As animal models continue to be paramount in understanding and treating cognitive impairment following TBI, the necessity of testing intervention strategies in clinically relevant settings cannot be ignored. This study used a unilateral parietal lobe controlled cortical impact (CCI) model of TBI and tested rats on a touchscreen-based associative learning task, Paired Associates Learning (PAL). In humans, PAL has been used to assess cognitive deficits in stimulus-location association in a multitude of disease states, including TBI. To date, the extent to which a rat model of TBI produces deficits in PAL has not yet been reported, although the usage of PAL will be important for understanding the clinical consequences of cognitive impairment post-injury and throughout intervention treatment. This study details the behavioral and histological consequences of the CCI injury model and closes a translational research gap between basic and clinical TBI research.

**HIGHLIGHTS:**

- PAL performance declines in a rat model of TBI.
- Response-driven bias in PAL becomes elevated after TBI.
- Inflammatory microglial response in the thalamus correlates with PAL deficit.

## 1. INTRODUCTION

Traumatic brain injury (TBI) in the U.S. continues to be a leading cause of disability for minors and young adults (CDC, 2019). Moreover, the rate of emergency department visits for TBI in individuals over 65 has increased by 78% from 2002 to 2017 (Cusimano et al. 2020). Acute effects of TBI can range from dizziness, nausea, headaches, and memory loss to motor coordination deficits, vision impairment, anxiety, irritability, and depression. Unfortunately, many individuals with TBI will also suffer from chronic neurocognitive symptoms that can lead to a decreased quality of life (McAlliser & Arciniegas 2002; Quinn et al. 2018). TBI occurs in two phases, the first of which is mechanical damage that occurs from the force of impact. The second phase of injury can occur in the days to weeks following initial insult and results from inadequately understood molecular cascades that often lead to the death of healthy and undamaged cells (Borgens & Liu-Snyder 2012). Due to the complexities of this secondary injury and the diverse array of cognitive dysfunction seen following TBI, animal models continue to be imperative in biomedical research aimed at understanding and treating TBI.

A promising avenue for the study of rat TBI models is the ability to monitor intervention outcomes in a clinically relevant setting. In humans, paired associates learning (PAL) is a touchscreen-based cognitive assessment within the Cambridge Neuropsychological Test Automated Battery (CANTAB) (Robbins et al. 1994, 1998; Barnett et al. 2016). Performance on PAL correlates with structural brain damage in individuals with TBI (Newcombe et al. 2016) and has been used in clinical trials testing the safety and efficacy of the cholinesterase inhibitor rivastigmine in TBI patients with memory impairment (Silver et al. 2006, 2009). Additionally, PAL was first developed and used in nonhuman primates (Weed et al. 1999; Taffe et al. 2002) and has since been modified for use in rats (Bussey et al. 2008; Talpos et al. 2009). The cross-species translational power of the touchscreen-based PAL task presents the ability to enhance the preclinical to clinical biomedical research pipeline. Despite the increasing imperative to conduct preclinical intervention testing that more closely reflects that seen in humans, no studies have yet detailed the extent to which a rat model of TBI leads to deficits in PAL performance that mirror those seen in human patients.

In rats, successful completion of the PAL task requires associative learning to accurately pair a stimulus (ex. flower, plane, or spider) with its correct location on the touchscreen (left, center, or right). While these object-place associations are known to require the hippocampus (HPC; Talpos et al. 2009; Kim et al. 2015; Langston et al. 2010; Yoon et al. 2012), the PAL task also likely involves structures relating to visual stimulus recognition, selective attention, executive control, and behavioral flexibility, all of which are commonly disrupted following TBI. The current study assessed performance on PAL after unilateral controlled cortical impact (CCI) to the parietal lobe of adult male and female rats. After the completion of cognitive testing, post-mortem histology was conducted to understand the extent of spared tissue in our model resulting from injury as well as the presence of glial fibrillary acid protein (GFAP) and ionized calcium-binding adapter molecule 1 (IBA1) in 12 regions of interest around the injury site. Furthermore, principal component analysis was performed to reduce the dimensionality of the dataset and inform how the various behavioral and histological measures were correlated with one another to aid future studies that use the PAL task in translational research for TBI treatment.

## 2. METHODS

### 2.1 Rats

A total of 16 adult (two months at time of arrival; n=8 male, n=8 female) Long Evans rats from Charles River were used in this study. Prior to completion of the study, three rats assigned to the sham condition and one animal assigned to the injury condition died from health complications unrelated to the CCI injury model. Each rat was housed individually in a standard Plexiglas cage and maintained on a 12-hour reversed light/dark cycle (lights off at 8:00 am). Testing was conducted exclusively in the dark phase of the cycle. Rats were given one week to habituate to the facility prior to behavioral testing or food restriction with *ad libitum* access to food (standard rat maintenance diet) and water. Throughout this habituation period, rats were handled daily. Upon the initiation of training, food was restricted to maintain the rats between 80-85% of their normal baseline body weight and had *ad libitum* access to water. Baseline weight was considered the weight at which an animal had an optimal body condition score of 3.0. Throughout the period of restricted feeding, rats were weighed daily, and body condition was assessed and recorded weekly to ensure a range of 2.5-3.5. The body condition score was assigned based on the presence of palpable fat deposits over the lumbar vertebrae and pelvic bones (Ullman-Culleré & Foltz, 1999; Hickman & Swan, 2010). Rats with a score under 2.5 were given additional food to promote weight gain. Food was provided *ad libitum* for the week following surgery. All procedures were conducted in accordance with the National Institutes of Health guidelines and were approved by the Institutional Animal Care and Use Committees at the University of Florida.

### 2.2 Surgery

After reaching a criterion performance on PAL, rats were pseudo-randomly selected, counterbalancing for sex, for either the TBI (n = 4 males/4 females), or sham (n = 4 males/4 females) condition. Initial anesthesia was induced by placing rats into a vented anesthesia chamber with 5% isoflurane. The head was shaved and then rats were placed in a stereotaxic frame and anesthesia was maintained with a nose cone at 1.5-2% isoflurane. Utilizing aseptic technique, a midline incision was made to expose the skull. A 7 mm diameter craniotomy was made, centered at AP -4.5 and ML + 4.5, targeting the rats’ right parietal lobe, as depicted in Figure 1A. A controlled cortical impact (CCI) was performed for rats in the experimental condition. The impact was delivered by a pneumatic piston (Model PCI3000 PinPoint Precision Cortical Impactor, Hatteras Instruments) containing a 5 mm diameter tip, which delivered a force at the rate of 4m/second, a 3mm compression and a 100ms dwell time Following the impact, the bone flap was replaced with bone wax, the incision was sutured, and rats were allowed to recover from anesthesia. Sham condition rats underwent similar surgical procedures but did not undergo a cortical impact. Rats had *ad libitum* access to food and water for seven days following surgery before returning to food restriction and PAL testing. Meloxicam injection (Alloxate; 5 mg/kg) was given before surgery and every 24 hours after surgery for 48 hours. After surgery, buprenorphine (Buprenex; 0.03 mg/kg) was administered every 8-12 hours for 24 hours, and rats received 0.5 ml of liquid antibiotic (Sulfamethoxazole and trimethoprim combination) with their food for 7 days. Rats were monitored post-surgery until time of sacrifice for any signs of distress. All procedures were conducted in accordance with the National Institutes of Health guidelines and were approved by the Institutional Animal Care and Use Committees at the University of Florida.

**Figure 1.**
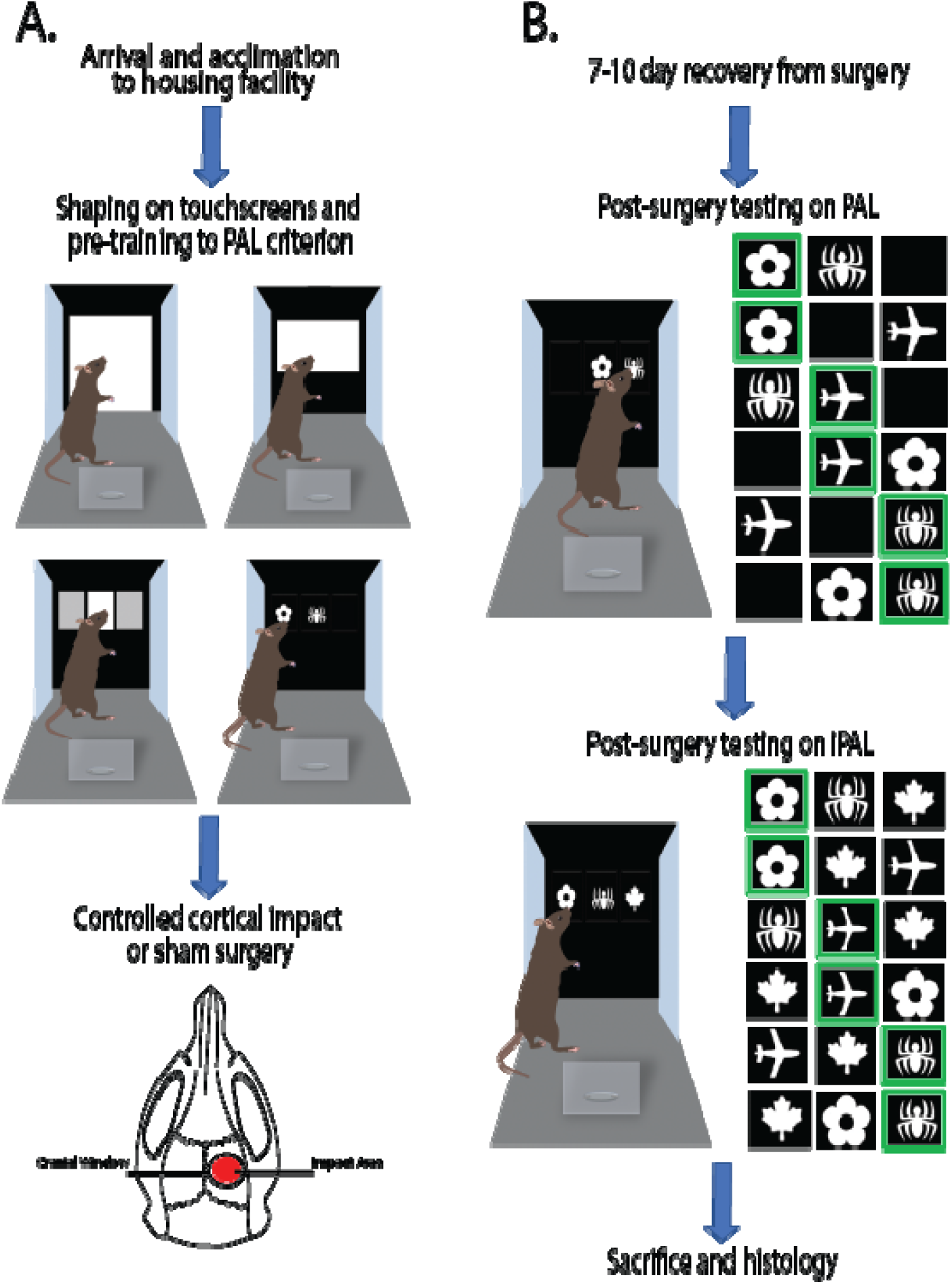
Schematic of experimental timeline. **A)** Pre-surgical experimental timeline. Rats were incrementally shaped to engage with the touchscreen chambers and pre-trained to reach PAL criterion. Following criterion achievement, rats were pseudo randomly selected for either CCI or sham surgery. **B)** Post-surgical experimental timeline. After a 7-10 day recovery from surgery, rodents were re-tested on PAL until criterion was met. Upon achievement of criterion, rats were tested for 12 days on iPAL. After completion of all behavior testing, animals were transcardially perfused for postmortem histological analysis.

### 2.3 PAL & iPAL

For an overview of the PAL and interference (i)PAL tasks as well as the shaping parameters used to accustom the rats to the touchscreen chambers, please see Smith, Zequeira et al. (2022). Rats were incrementally shaped to engage with the touchscreen apparatus before being tested on PAL. The testing was conducted using standard Bussey-Saksida rat touchscreen chambers (Lafayette Instruments, Lafayette, IN) that were housed within soundproof boxes. Engaging with the touchscreen apparatus requires a nose-poke on the correct stimulus to elicit a liquid reward of Ensure nutritional shake. After the reward is collected, the next trial begins after a 10-second inter-trial interval. If the nose-poke occurs on the incorrect stimulus, no reward is received, and a 10-second timeout period occurs, during which the house light turns on and another trial cannot be initiated. Rats were trained to complete 90 trials in a daily session. The PAL task incorporates 3 possible stimuli and 3 possible locations on the touchscreen. During a trial, two stimuli are presented at the same time; one stimulus is presented in the correct location and the other in the incorrect location. The unused, third location of a given trial is a blank panel and any engagement with this blank panel does not elicit any change in the environment (no reward or house light). Each of the three stimuli are assigned to a correct location that is unique to the stimuli, resulting in six different trial types, shown in figure 1B. Each PAL and iPAL session lasted for a maximum of 45 minutes or concluded upon completion of 90 trials.

Rats were tested on the PAL task until they reached a criterion performance of 83.33% correct for two consecutive testing days. After completion of criterion on PAL, rats underwent CCI surgery (see above). Following recovery, rats were re-tested on PAL until criterion performance was again met. Once criterion on PAL was met post-surgery, rats then moved on to complete 12 days of iPAL testing before sacrifice and postmortem histology. The iPAL task is a modification of PAL in which a novel interference image (a white maple leaf) is presented in the blank panel as a distractor. Selection of the novel stimulus does not elicit a reward, but rather is counted as an incorrect trial and followed by a 10-second timeout with the house light on. Figure 1 depicts the six possible PAL and iPAL trial types, a schematic of a rat completing a trial in the touchscreen environment, and a timeline of the experimental procedures.

### 2.4 Histological Analysis

Rats were transcardially perfused with 4% paraformaldehyde in phosphate buffered saline. Brains were post-fixed for 24 hours and sliced into 50-micron thick sections using a vibratome. Each microscopy slide was plated with sequential slices approximately 350 microns apart. Sections were Nissl stained (1% Cresyl Violet) for lesion volume analysis. Tissue was also stained for the presence of GFAP and IBA1 epitopes. GFAP is an intermediate filament protein expressed by astrocytes in the central nervous system that has been viewed as an index of gliosis and correlate of neural damage (Finch 2003; Hausmann, 2003). IBA1 is a calcium-binding protein in microglia and macrophages that is involved in phagocytosis in activated microglia (Ohsawa et al. 2004). As microglia move to an activated state after exposure to pathogen and damage-associated molecular patterns (Jurga, Paleczna, & Kuter 2020), this protein is associated with pro-inflammatory processes.

For the GFAP and IBA1 immunohistochemistry, primary antibody was used at [1:10,000] for GFAP (GA5 mAb #3670; Cell Signaling Technology) and at a [1:10,000] for IBA1 (ab107159; Abcam). Secondary antibody was used at [1:2,000] for GFAP (rabbit anti-mouse biotinylated; ab107159; Abcam) and [1:2,000] for IBA1 (rabbit anti-goat biotinylated; ab64257). Sections were mounted on Superfrost Plus slides, air dried, and placed in 0.1 M Tris buffer, pH 7.6 (Tris). Slides were then immersed in 85– 87°C Tris for 1 minute, washed in Tris, and placed in Tris containing 0.25% bovine serum albumin (BSA; fraction V; Sigma) and 0.1% Triton X-100. After primary antibody incubation, slides were washed in Tris-BSA-Triton X buffer, pH 7.6 (2-5 minutes minimum). Slides were incubated in secondary antibody solution in TrisBSA-Triton X buffer for 2 hours, washed in the same buffer, and then incubated for 2 hours in avidin-biotinHRP complex (PK-6100; Vector Laboratories ABC Elite Kit diluted [1:1,000] in TrisBSA-Triton X buffer). Slides were then washed in Tris (3 5 minutes minimum) and incubated in a hydrogen peroxide-generating 3-3’-diaminobenzidine (DAB) solution (100 ml Tris containing 50 mg DAB, 40 mg ammonium chloride, 0.3 mg glucose oxidase, and 200 mg β-D-glucose). After incubation in DAB solution (20 –30 minutes), slides were rinsed in Tris, dehydrated in graded ethanols and xylene, and coverslipped with Permount. Brightfield images were acquired on a Keyence BZ-X800 microscope (Keyence Corporation of America, Itasca, IL).

For semi quantitative analysis of the GFAP & IBA1 stains, 12 regions of interest were identified, depicted in Figure 2A. Five regions of interest were defined a priori within the hippocampus due to the high degree of involvement of this structure required for the cognitive testing used in this study: proximal CA1 (p.ca1), CA1, subiculum (sub), CA3, and CA3c area (CA3c). Seven additional thalamic regions were later identified for inclusion: lateral posterior nucleus - mediorostral part (lpmr), lateral dorsal nucleus - ventrolateral part (ldvl), posterior nuclear group (po), ventral posteromedial nucleus (vpm), ventral posterior lateral nucleus (vpl), mediodorsal nucleus – medial part (mdm), and mediodorsal nucleus – lateral part (mdl). The optical density in these regions of interest was quantified across three slices for each animal in each hemisphere. Images of the DAB-stained sections were first captured at 10X magnification. In FIJI (NIH, Bethesda MD), the images were color deconvoluted and converted into 8-bit grayscale. A pre-designed set of ROIs with standard sizes were utilized. For each slice, experimenters blinded to condition identity adjusted the ROI boxes to best match the tissue’s own anatomical features. After calibrating the Measure function to an optical density standardization image, an optical density curve was created for use in measuring each ROI. Within-slice ROI measurements were normalized by the mean gray value of that slice to account for variability in tissue darkness.

**Figure 2.**
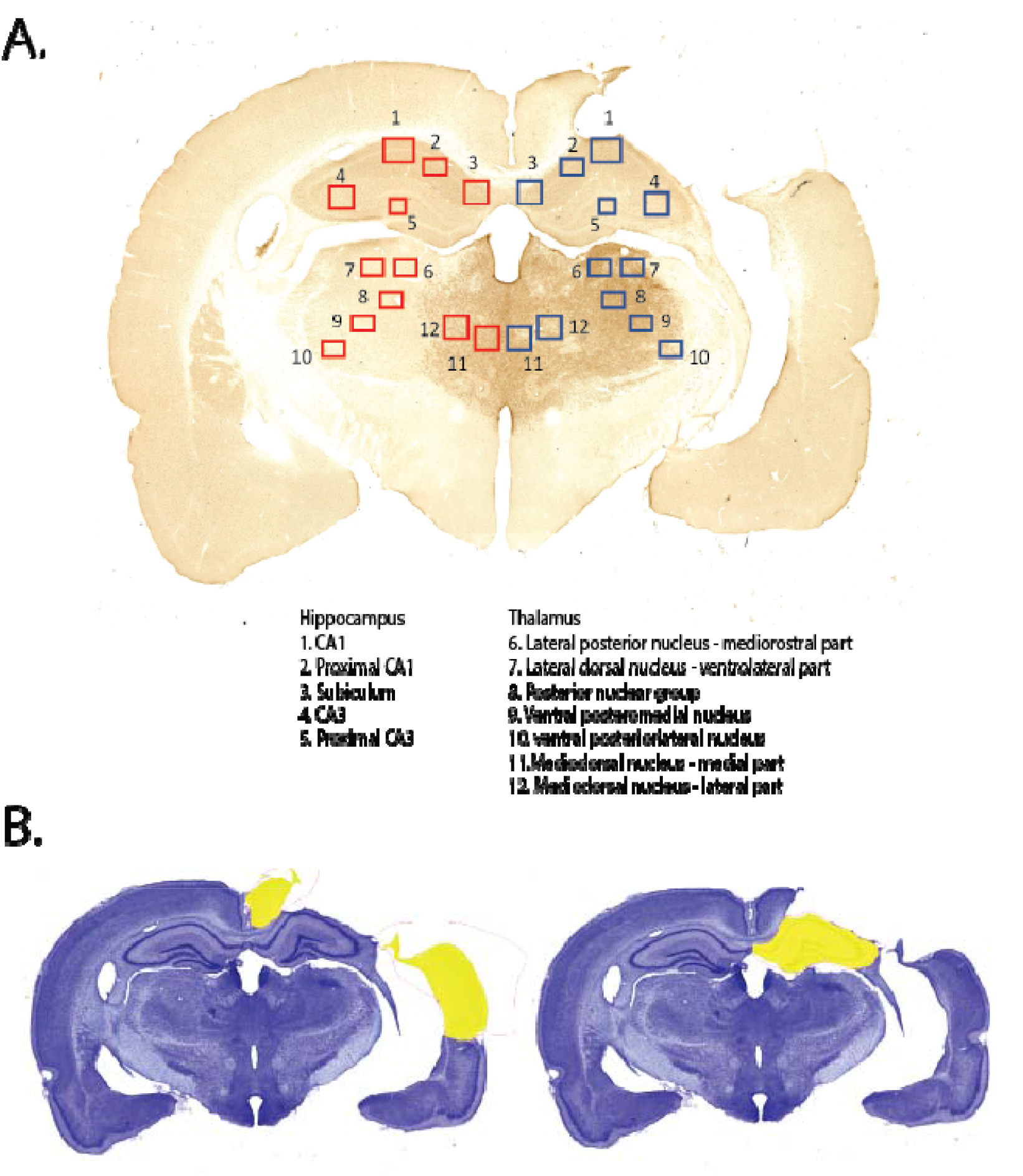
Depiction of parameters for postmortem histological analysis. **A)** IBA1 stained section for a CCI rat with the 12 regions of interest identified for analysis superimposed on the ipsilateral (dark blue) and contralateral (red) hemispheres. **B)** Images of nissl-stained sections from a CCI rat highlighting spared lesion volume tracings (yellow) for the cortex (left) and hippocampus (right).

For spared tissue volume analysis, eight representative Nissl-stained sections across the extent of injury were imaged on a Keyence (BZ-X800) at 2X magnification. Using the Hybrid Cell Count module in the BZ-X800 Analyzer software, the area (µm^2^) of each hemisphere was recorded, as well as that of the ipsilateral hippocampus and cortex (dorsal to rhinal sulcus), the latter two being hand-traced by a blinded experimenter as depicted in Figure 2B. Ipsilateral hemisphere, hippocampal, and cortical measurements were normalized by the slices’ contralateral hemisphere to mitigate between subject variability in brain size.

### 2.5 Analytical & Statistical Parameters

All trial-by-trial analyses were conducted using custom written code (MATLAB, MathWorks Inc, R2021A) that is available upon request. For an in-depth overview of how parameters such as bias and strategy were calculated, please see Smith, Zequeira et al. (2022). Briefly, all bias was calculated as

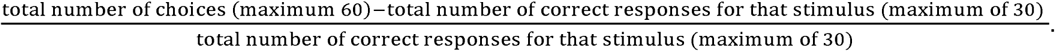

Bias ranges from 0 (no bias) to 2 (absolute bias). Win-stay and lose-shift strategies were calculated as total number of times a strategy was used / total number of times a strategy was possible. For each of the 4 possible strategies (stimulus win-stay, stimulus lose-shift, location win-stay, and location lose-shift), with bias ranging from 0 (no strategical use) to 1 (maximal strategical use). Statistical analyses were performed using the statistical software R.3.6.3. Whenever possible, we elected to fit a mixed effects linear model using the R package “lme4” (version 1.1-28; https://github.com/lme4/lme4/) to assess statistical significance due to the limitations of ANOVA in considering variability between subjects (Barr et al. 2013; Brown, 2021), especially for repeated measures designs in which both within and between subjects’ variables are present. P-values < 0.05 were considered statistically significant, and ‘trends toward significance’ statements were made when the p-value was less than 0.2 accompanied by a noticeable effect size in the graphical representation of the data. Bonferroni corrections were used for multiple comparisons. Reporting of the linear model effects in this manuscript follow Satterthwaite’s method (Giesbrecht & Burns 1985; Kuznetsova, Brockhoff, & Christensen 2017). When appropriate, behavioral data were analyzed by t-tests with correction for unequal variances if necessary. The choice of statistical test was based on assumptions of normality, assessed with the Shapiro-Wilk test and with Levene’s test.

## 3. RESULTS

### 3.1 PAL Performance

To assess post-surgical PAL performance, a linear mixed effects model (LMEM) was fit to predict percent correct with both condition (sham or injury) and day as fixed effects. The model also included the individual rats as a random effect. Although some rats completed more than 12 days of PAL testing, all rats only completed 12 days of iPAL testing. For consistency of reporting, all analyses in this section are across 12 days of PAL testing. The model’s power was substantial with a conditional R^2^ value of 0.63 and a marginal R^2^ of 0.22. The model’s intercept was 78.55 (95% CI [71.63, 85.47], T_(138)_ = 22.44, p < .001). This model found significant main effects of condition (F_(1,15.45)_ = 7.157 p = 0.02) and day (F_(1,130)_ = 4.059, p = 0.046), as well as a significant day by condition interaction (F_(1,130)_ = 6.067, p = 0.015). Figure 3A shows the average percent correct of each condition on the day before surgery as well as the first 12 days of testing after recovery from surgery. When sex was added to the model, it did not explain any additional variance F_(1,11.44)_ = 0.017, p = 0.898, standardized β = -0.02). Figure 3B visualizes the results of post hoc un-paired t-tests. Before surgery, both conditions performed at similar levels (T_(10)_ = 0.292, p = 0.775). Post-surgery, a five-day average of performance found that the conditions were statistically different (T_(10)_ = 2.378, p = 0.039). Furthermore, Figure 3C shows the cumulative error (number incorrect / total trials) prior to reaching criterion on PAL post-surgery. The sham condition had an average of 223.2 ± 44.94 S.E.M. errors across testing. The injury condition had 317.2 ± 27.43 S.E.M. An unpaired t-test did not reach statistical significance (T_(10)_ = 1.894, p = 0.088), although there was a trend for the CCI treated rats to make more errors. Overall, the data presented in figure 3A-C indicates that rats that received CCI were less accurate on PAL following surgery compared to sham rats and required more time to return to criterion performance.

**Figure 3.**
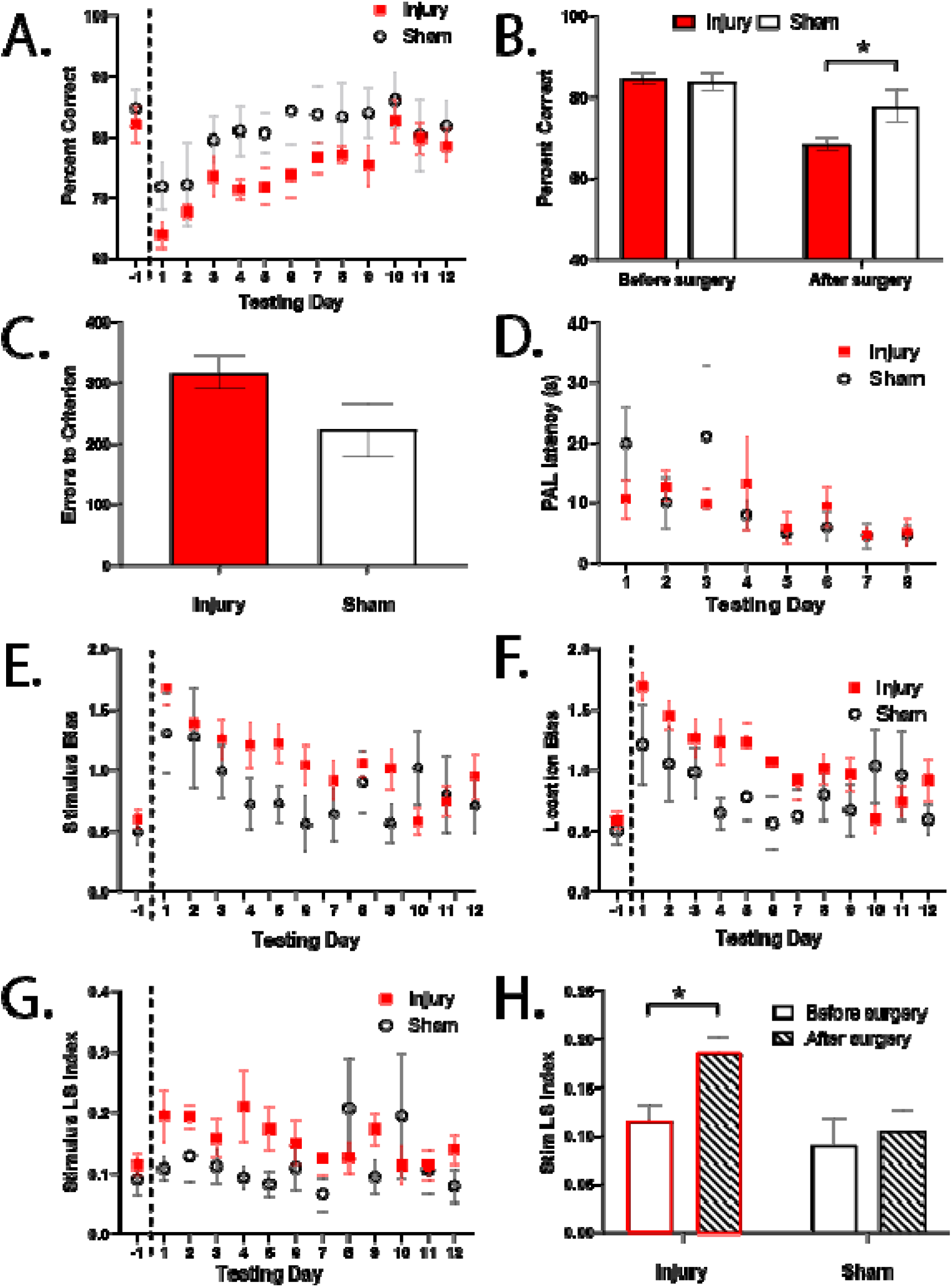
CCI resulted in PAL performance deficits compared to sham condition. **A)** Post-surgical PAL performance was significantly affected by injury condition and day of testing, as well an interaction between the two as measured by percent correct (p<0.05 for all measures). **B)** Before surgery, bot conditions performed comparably on PAL. After surgery, injury condition had signficantly reduced percent correct (p<0.05). **C)** After surgery, injured rats trended toward having more errors on PAL prior to reaching criterio (p<0.1). **D)** There was no effect of injury condition on PAL trial latency after surgery. **E)** Post-surgical stimulus bias on PAL was signficantly affected by injury condition (p<0.05) and an interaction between day and injury conditio (p<0.01). **F)** Post-surgical location bias on PAL was significantly affected by injury condition (p<0.01) as well as an interaction between day and injury condition (p<0.001). **G)** Post-surgical stimulus lose-shift index on PAL was significantly affected by injury condition (p<0.01) as well as an interaction between day and injury condition (p<0.05). **H)** Injury condition showed a signficant increase in stimulus lose-shift index from pre to post-surgery (p<0.05), but sham condition did not.

To rule out whether differences in motor coordination or motivation may have affected performance between the conditions, we fit a LMEM for trial latency (time between the start of a new trial and when a decision is made, in seconds). Figure 3D shows the average latency for each condition across the first 8 days of testing post injury. This model found a significant main effect of day (F_(1, 124.9)_ = 13.89, p = 0.0003) but not condition (F_(1,36.87)_ = 0.373, p = 0.545). Furthermore, the day by condition interaction was also nonsignificant (F_(1,124.7)_ = 0.258, p = 0.613). These results suggest that there was no difference in trial latency between the conditions, making it unlikely that any potential motor deficits in the injury condition affected performance on PAL. Furthermore, the finding that both conditions’ latencies increased across testing day indicates that motivation increased with the learning and familiarity of the task regardless of injury condition, which was expected with similar levels of motivation across conditions.

In addition to the accuracy and latency measures detailed above, the PAL task presents the ability to run a trial-by-trial analysis of performance measures across a 90-trial session. For these trial-by-trial analyses, we first looked at trial blocks over the course of a session across testing days to understand if the two conditions of rats diverged in terms of performance in earlier vs. later trials in a session. This was achieved through blocking the daily 90-trial sessions into two blocks of 45 trials and taking the difference between the percent correct values in these epochs. We then fit a LMEM. This model found no significant effects of day, condition, or day by condition interaction (day: F_(1, 128.2)_ = 0.944, p = 0.333, condition: F_(1,46.84)_ = 0.278, p = 0.601, day by condition: F_(1,128.1)_ = 0.0054, p = 0.941) suggesting that both conditions had similar trajectories of performance in early and late trials. The addition of sex to the model did not account for any additional variance seen in block performance (p > 0.05, β < 0.1). These null results are not depicted in graphical format.

The next trial-by-trial analysis we conducted was a bias index for feature parameter selections during a testing session. Bias is the tendency for a rat to default to a response-driven behavior, choosing a feature parameter repeatedly. In the case of PAL, bias can be towards a certain location (for example, choosing the left panel repeatedly even when incorrect), or a certain stimulus (for example, choosing flower repeatedly). Figure 3E and 3F shows the average bias by condition on the day before surgery and across 12 days of testing post-surgery for stimulus and location features, respectively. For stimulus bias, LMEM found a significant main effect of condition (F_(1,19.29)_ = 6.415, p = 0.020) as well as a significant condition by day interaction (F_(1,130)_ = 7.380, p = 0.008). Day alone was non-significant (F_(1,130)_ = 1.316, p = 0.2534). Similarly, for location bias, LMEM found a significant main effect of condition (F_(1,17.87)_ = 11.08, p = 0.004) and condition by day interaction (F_(1,130)_ = 15.30, p = 0.00015), while day alone was non-significant (F_(1,130)_ = 0.547, p = 0.461). These results suggest that the CCI-injured rats exhibited higher response biases for both stimulus and location features across testing than sham rats, which likely contributed to their overall performance deficit as this bias has been shown previously to negatively correlate with percent correct (Smith, Zequeira et al. 2022). Moreover, when the difference between stimulus and location bias was fit, there was a significant main effect of condition (F_(1,140)_ = 5.130, p = 0.025), suggesting that in the injured condition, biases varied more between one another.

In addition to having a response-driven bias, rats could also adopt an outcome-based strategy. On a given trial, they could either select a previously rewarded feature (win-stay) or switch from a previously unrewarded feature (lose-shift). For example, if a rat incorrectly chooses the flower in a trial and is then immediately given another trial where flower is now correct, if the rat then does not choose flower, this can be considered a stimulus lose-shift strategy. A strategy usage index was calculated for four outcome-based heuristics in PAL before and after surgery: stimulus win-stay, location win-stay, stimulus lose-shift, and location lose-shift. On the day before surgery, all strategy usage was the same for both surgery conditions (stim ws: T_(10)_ = 0.078, p = 0.9394; loc ws: T_(10)_ = 0.128, p = 0.901; stim ls: T_(10)_ = 0.762, p = 0.464; loc ls: T_(10)_ = 0.261, p = 0.799). After injury, stimulus win-stay, location win-stay, & location lose-shift did not significantly differ by condition (F_(1,83.763)_ = 3.794, p = 0.055; F_(1, 68.039)_ =1.991 p =0.163; F_(1,140)_ = 1.250, p = 0.2655). Stimulus lose-shift, however, had a significant condition (F_(1,45.947)_ = 7.417, pN= 0.009) and condition by day (F_(1,130)_ = 4.593, p = 0.034) interaction. The effect of day alone did not reach significance (F_(1,130)_ = 0.528, p = 0.468). Furthermore, from pre to post surgery (five-day average), stimulus lose-shift did not significantly differ in the sham condition (T_(8)_ = 0.451, p = 0.664) but did in the injury (T_(12)_ = 2.955, p = 0.012). Figure 3G and 3H illustrates this stimulus lose-shift data. Taken together, these results suggest that the injured condition had an increase in stimulus lose-shift strategy usage on PAL as a result of TBI. Sex did not affect any strategy usage, either pre or post injury, and did not interact with condition (p>0.05, β < 0.1 for all measures).

### 3.2 iPAL Performance

After reaching PAL criterion post-surgery, both conditions were then tested on iPAL, a modification of PAL created to assess vulnerability to irrelevant distractors. Figure 4A shows the average percent correct across 12 testing days. A LMEM fitting percent correct on iPAL with condition and day as fixed effects and subject as a random effect did not find the conditions statistically different (condition: F_(1,12.35)_ = 0.975, p = 0.342, day: F_(1,119)_ = 0.596, p = 0.442, interaction: F_(1,119)_ = 1.290, p = 0.258.) Additionally, adding sex to the model did not account for any additional variance (F_(1, 9.058)_ = 0.351, p = 0.568, standardized β = 0.06). Figure 4B shows the cumulative error that occurred across 12 days of iPAL testing. The sham condition had an average of 331.3 ± 53.65 S.E.M errors. The injury condition had an average of 422.4 ± 35.01 S.E.M. An unpaired t-test did not find these conditions statistically different (T_(9)_ = 1.469, p = 0.176), but the injured condition showed a trend toward more errors.

**Figure 4.**
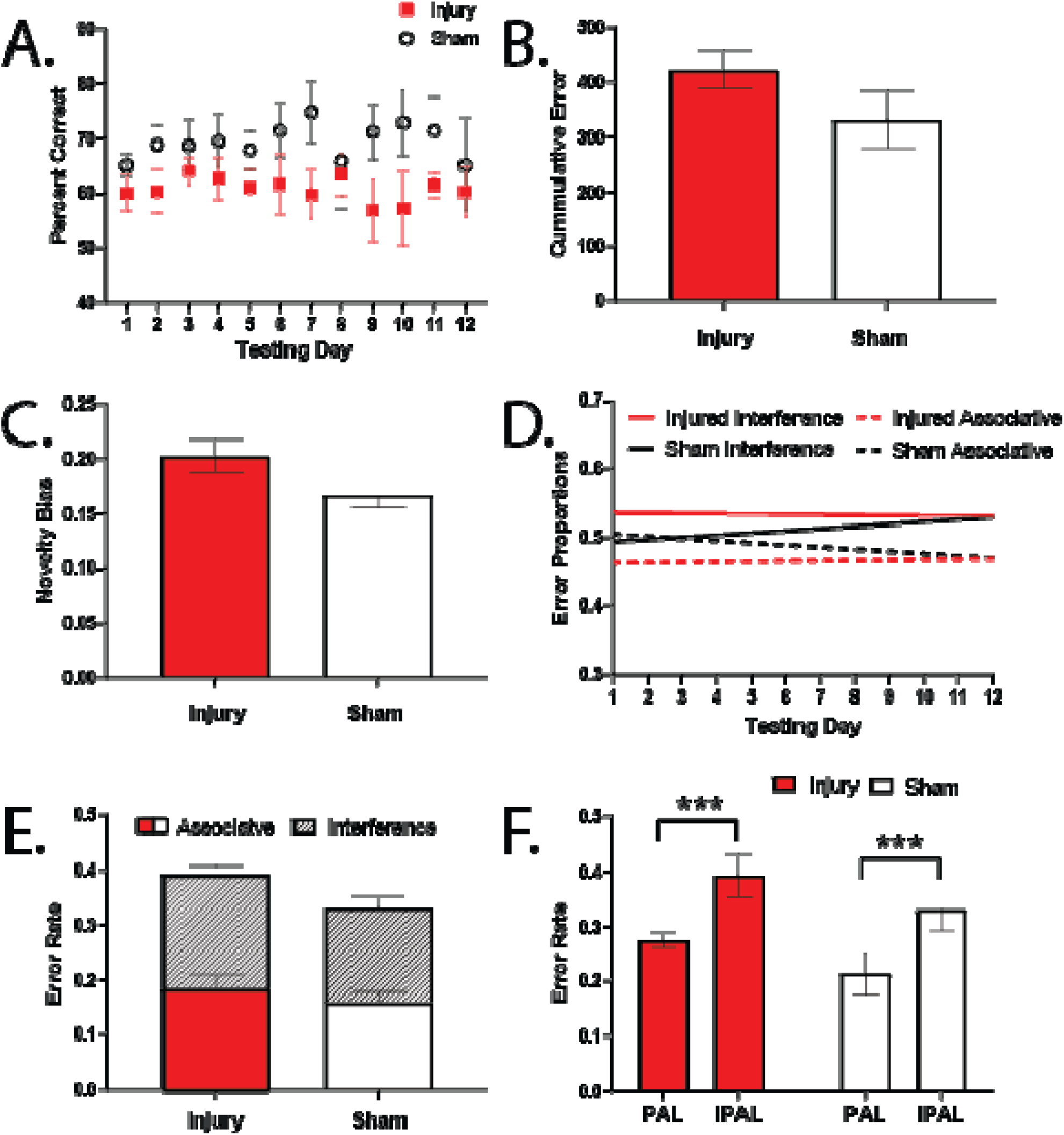
CCI did not result in statistically significant iPAL performance deficits compared to sham condition. **A)** Post-surgical iPAL performance was not significantly affected by injury condition, day, or condition by day interaction (p>0.05). **B)** Injury condition showed a trend toward more errors on iPAL (p<0.2) with modest effect size. **C)** Injury condition showed a trend toward more interfence errors as measured by novelty bias (maple leaf touches / total trials) (p<0.2). **D & E)** Both injury and sham conditions showed similar proportions of error type across iPAL testing (p>0.05). **F)** Both injury and sham conditions showed increased error rate from PAL to iPAL (p<0.0001).

To quantify susceptibility to the maple leaf distractor, we evaluated novelty bias by calculating the ratio of: [number of maple leaf touches in a session / number of trials in a session]. The average across the 12-testing days of iPAL for each condition is depicted in Figure 4C. The difference in the conditions again trended toward significance (t(8) = 1.659, p = 0.136). In iPAL, two types of errors can be made: interference error (error made by selecting the irrelevant distractor, i.e., novelty bias) and associative error (error unrelated to the distractor, the only error seen in standard PAL). Figure 4D shows the linear regression of these two types of errors as a proportion of the total error made on iPAL across the 12 testing days. A LMEM was fit for the difference between the two proportions across the 12 testing days with condition and day as fixed effects and animal as a random effect. This model found no significant effects of condition, (F_(1, 58.14)_ = 1.255, p = 0.267), day (F_(1, 119)_ = 0.961, p = 0.329), or condition by day interaction (F_(1, 119)_ = 0.645, p = 0.423). These results suggest that each condition experienced a similar proportion of error type on iPAL. Visualized another way, Figure 4E shows the error proportions as a function of the overall error rate on iPAL as an average across the 12 testing days. It is apparent that each error type makes up approximately half of the total error for both conditions. Finally, Figure 4F shows the error rates for both conditions averaged across the first 12 days of testing on PAL versus iPAL. To compare the error rates between PAL and iPAL, a 2-way ANOVA with PAL and iPAL error rates as a within subjects’ factor and injury condition as a between subjects’ factor revealed a main effect of PAL versus iPAL (F_(1,9)_ = 42.76, p = 0.0001) but no effect of condition (F_(1,9)_ = 2.689, p = 0.136) nor interaction between condition and PAL/iPAL error rates (F_(1,9)_ = 0.465).

Taken together, these results suggest that the overall error rate for both conditions on iPAL was due to a similar proportion of associative and interference errors, and that both conditions exhibited additional error on iPAL that was interference related. While the iPAL measures presented here were not statistically significant, Figures 4A and 4B show modest effect sizes with large S.E.M.s in both conditions. Furthermore, any trend toward impairment on iPAL in the injury condition can be interpreted as a continuation of the associative learning deficit that was seen during the standard version of PAL (Figure 3), rather than an increase in vulnerability to irrelevant distractors.

### 3.4 Spared Tissue Volume

During tissue processing, significant variability in lesion volume was qualitatively observed. While some CCI-injured rats had large lesions, others had significant tissue swelling around the injury region. Figure 5A-D shows Nissl-stained tissue sections from four rats that were representative of the different patterns we observed in the injury (Figure 5A and 5B) and sham (Figure 5C and 5D) conditions. Of the seven injured rats, one was characterized by inflammation (representative animal shown in Figure 5A), three were characterized by having a ‘hole’ (representative animal shown in Figure 5B), and three were characterized as some combination of ‘inflammation’ and ‘hole’. There was no variability in the sham condition (Figure 5C and 5D). Three different quantitative measurements of spared tissue around the injury site were taken for each rat as outlined in section 2.3. With subject variability accounted for in the LMEM, we found significant effects of condition (F_(1,10)_ = 8.74, p = 0.014), spared tissue volume measurement (F_(2,20)_ = 1021, p < 0.0001), and condition by volume measurement interaction (_F(2,20)_ = 7.863, p = 0.003) in the spared tissue assessments. The addition of sex to the model did not explain any additional variance (p>0.05, β < 0.1 for all measures). Figure 5E shows the quantifications of spared tissue volume. Overall spared tissue volume (normalized to contralateral hemisphere) was significant (T_(10)_ = 3.027, p = 0.013), as well as cortical spared volume (T_(10)_ = 3.019, p = 0.013), but hippocampal spared volume was insignificant (T_(10)_ = 0.249, p = 0.808). Furthermore, an F-statistic of variance trended toward significance (F_(6,4)_ = 0.197) for the overall spared tissue volume.

**Figure 5.**
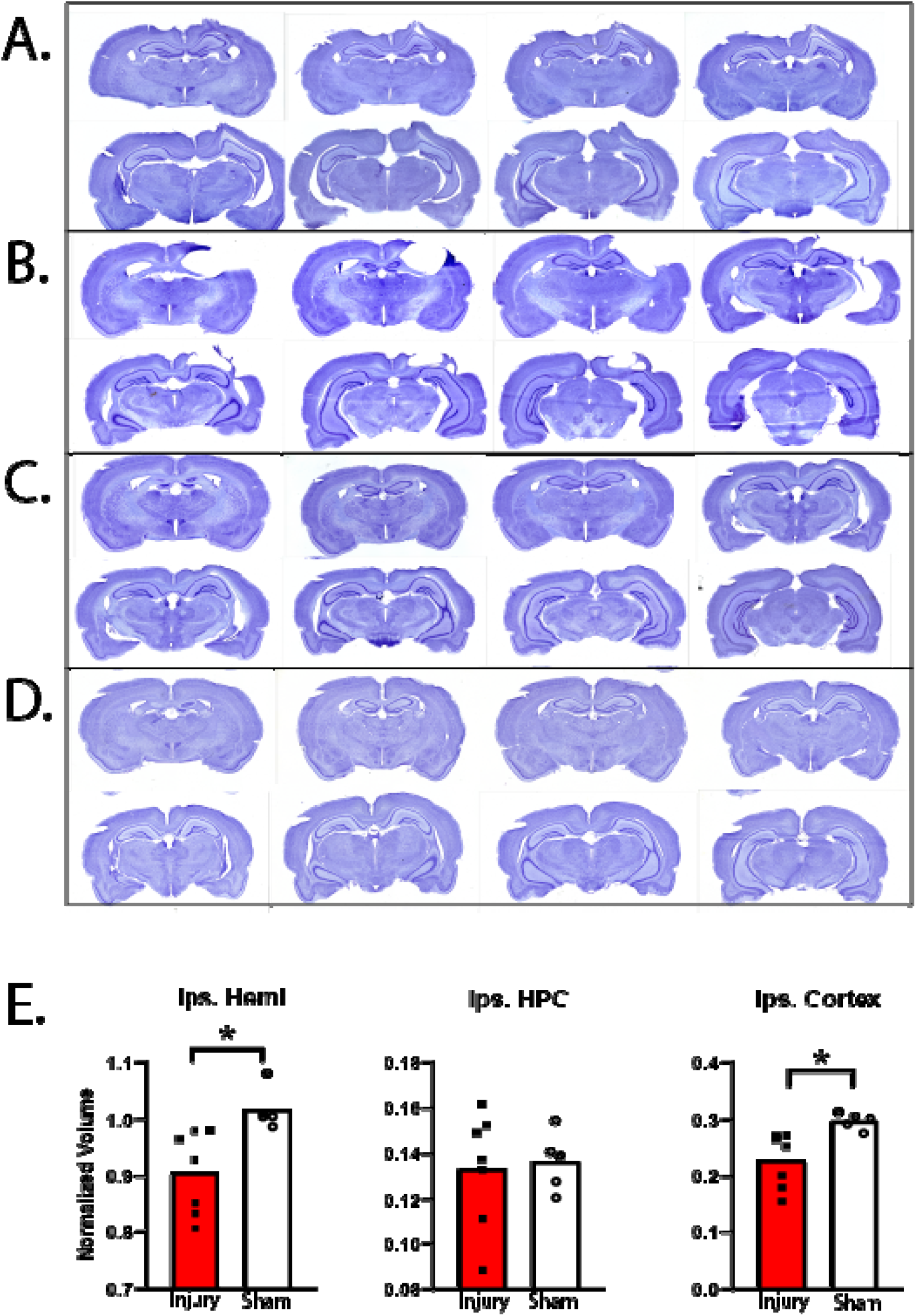
CCI resulted in greater tissue loss compared to sham condition as measured by spared tissue volume. **A-D)** Nissl-stained sections across extent of injury for two different CCI rats (A,B) and two different sham rats (C,D). Qualitative variability in lesion volume between CCI rats can be seen by the different patterns of injury exhibited in the sections of rat A versus rat B. **E)** CCI rats had lower spared tissue volume compared to sham in the hemisphere ipsilateral to injury site (left; p<0.01). Separating the measurements into hippocampal (middle) or cortical (right) spared volume, injured animals only differed from sham in spared cortical volume (p<0.05).

### 3.5 Semiquantitative immunochemistry

Immunochemistry was performed on the tissue to assess the presence of GFAP and IBA1 antibody signal in the predefined 12 regions of interest, both ipsilateral and contralateral to the side of injury. Figure 6A-C shows microcopy images taken at 10X of the GFAP staining in three injured (A-C) and one sham (D) rat. Figure 6E shows 20X magnification images of the regions outlined in panel A. Figure 6F, 6G, and 6H shows the quantification of GFAP optical density between conditions in the ipsilateral ROIS, between conditions in the contralateral ROIs, and ipsilateral versus contralateral side within the injury condition, respectively. LMEM was fit with condition, regions, and side as fixed effects, and subject as a random effect. This model revealed significant main effects of condition (F_(1,10.00)_ = 7.336, p = 0.022) and region (F_(11,229.00)_ = 11.016, p < 0.0001). Furthermore, there were significant interaction effects of condition by region (F_(11,229.1)_ = 1.888, p = 0.042), condition by side (F_(1,229.006)_ = 29.773, p < 0.0001), and condition by region by side (F_(11, 229.00)_ = 1.832, p = 0.050. When sex was added to the model, it did not account for any additional variance (p = 0.322, β = 0.07).

**Figure 6.**
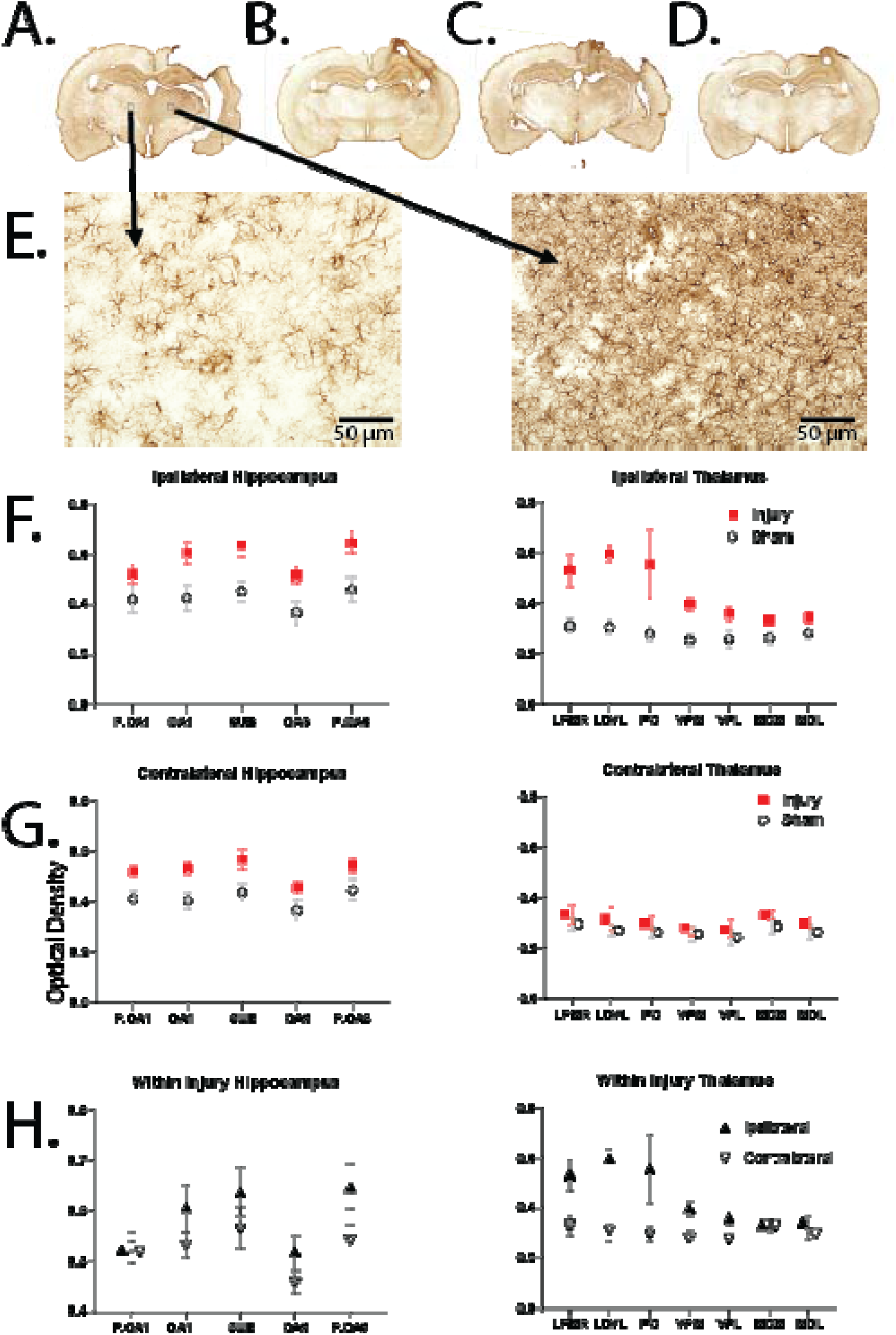
CCI resulted in elevated GFAP staining as measured by optical density compared to sham condition. **A-D)** 10X stitched images of GFAP stained sections for 3 different CCI rats (A-C) and 1 sham rat (D). **E)** 20X images taken from the regions defined in panel A. F) CCI resulted in greater optical density of GFAP in the side ipsilateral to the injury site compared to sham. Both hippocampus (left) and thalamus (right) had significant effects of condition (p<0.0001). G) CCI resulted in greater optical density of GFAP in the side contralateral to injury site compared to sham in the hippocampus only (p<0.01). F) Within the CCI condition, rats had greater optical density of GFAP on the side ipsilateral to injury for both hippocampus (left; p < 0.01) and thalamus (right; p < 0.0001) ROIs.

Further two-way ANOVAs were ran for GFAP optical density. Both ipsilateral hippocampal and thalamic ROIs had a significant main effect of condition (hippocampus: F_(1,50)_ = 31.64, p < 0.0001; thalamus: F_(1,70)_ = 32.85, p < 0.0001). Ipsilateral thalamus ROIs also had a significant effect region (F_(6,70)_ = 2.944, p = 0.0128), but the interaction between condition and region was nonsignificant (p = 0.173). Ipsilateral hippocampal ROIs had no other significant effects (p>0.05). Both contralateral hippocampal and thalamic ROIs had a significant main effect of condition (hippocampus: F_(1,50)_ = 27.71, p < 0.0001; thalamus: F_(1,70)_ = 4.265, p = 0.043), but no other effects (p>0.05). Finally, within our injury condition, ANOVA revealed significant main effects of both side and region for hippocampus and thalamus (hpc: F_(1,60)_ = 7.757, p = 0.007, F_(4,60)_ = 3.797, p = 0.008; thal: F_(1,84)_ = 29.67, p < 0.0001, F_(6,84)_ = 3.101, p = 0.009). The interaction between side and region in the thalamic ROIs reached significance F_(6,84)_ = 2.447, p = 0.031, but becomes nonsignificant when adjusted for multiple comparisons.

Figure 7A-C shows microcopy images taken at 10X of the IBA1 staining in three injured (A-C) and one sham (D) rat. Figure 7E shows 20X magnification images of the regions outlined in panel A. Figure 7F, 7G, and 7H shows the quantification of IBA1 optical density between conditions in the ipsilateral ROIS, between conditions in the contralateral ROIs, and ipsilateral versus contralateral side within the injury condition, respectively. LMEM was fit with condition, regions, and side as fixed effects, and subject as a random effect. This model revealed a significant main effect of region (F_(11,230)_ = 33.89, p < 0.0001). Furthermore, there were significant interactions of condition by region (F_(11,230)_ = 2.267, p = 0.012), and condition by side (F_(1,230)_ = 10.76, p = 0.001.) Unexpectedly, when sex was added to the model, there was a significant main effect of sex (F_(1,8)_ = 6.154, p = 0.03807), as well as an interaction between condition and sex (F_(1,8)_ = 10.6994, p = 0.01134). While the main effect does not reach significance after correction, the interaction does. However, because we are not sufficiently powered to detect sex differences (sham females n=2), we cannot make any conclusionary remarks. Nonetheless, this may be an interesting avenue to pursue in the future regarding sex differences in IBA1 expression following TBI.

**Figure 7.**
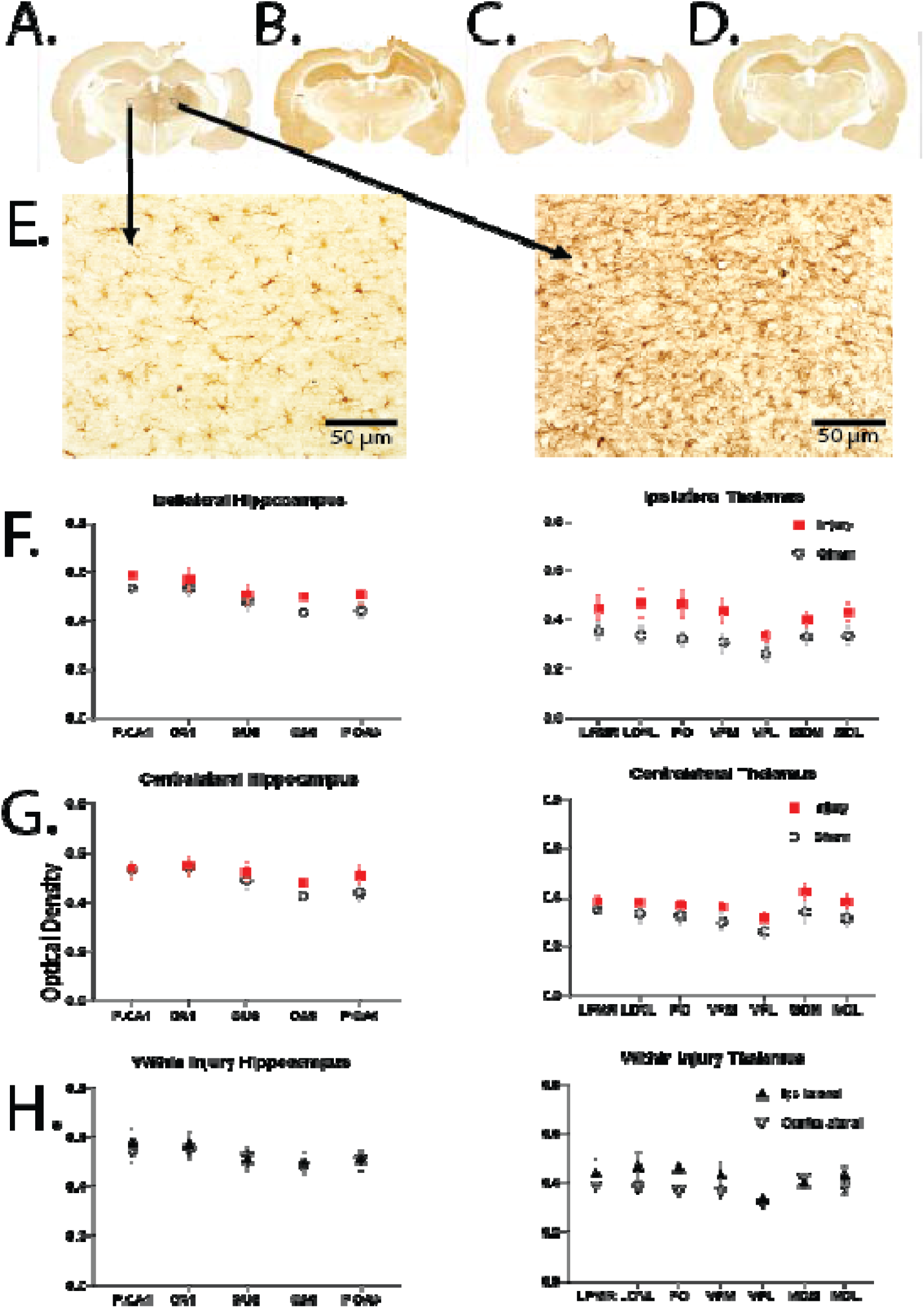
CCI resulted in elevated IBA1 staining as measured by optical density compared to sham condition. **A-D)** 10X stitched images of IBA1 stained sections for 3 different CCI rats (A-C) and 1 sham rat (D). **E)** 20X images taken from the regions defined in panel A. F) CCI resulted in greater optical density of IBA1 in the side ipsilateral to injury site only in the thalamic ROIS (p < 0.0001). G) CCI resulted in greater optical density of IBA1 in the side contralateral to injury site only in the thalamic ROIs (p < 0.001). H) Within the CCI condition, rats had a greater optical density of IBA1 on the side ipsilateral to injury for only the thalamic ROIs (p < 0.01).

Further two-way ANOVAs were ran for IBA1 optical density. Ipsilateral thalamic, but not ipsilateral hippocampal, ROIs had a significant main effect of condition (F_(1,70)_ = 19.54, p < 0.0001). No other effects were found for ipsilateral ROIs between conditions. (p>0.05). Furthermore, contralateral thalamic, but not contralateral hippocampal, ROIs had a significant main effect of condition (F_(1,70)_ = 13.90, p = 0.0004). Ipsilateral thalamic ROIs also had a main effect of region (F_(6,70)_ = 2.237, p = 0.049), but the interaction between condition and region was nonsignificant when adjusted for multiple comparisons (F_(6,70)_ = 2.237, p = 0.049). There were no other effects (p>0.05) for contralateral ROIs between conditions. Within our injury condition, ANOVA did not reveal any significant effects of contralateral/ipsilateral side, region, or their interaction for the hippocampal ROIs (p > 0.05 for all measures). For the thalamic ROIs, ANOVA revealed only a significant main effect of side (F_(1,84)_=7.109, p = 0.009).

### 3.6 Principal Component Analysis

Principal component analysis (PCA) was used as a dimensionality reduction technique on the variables obtained in this study to simplify correlational results and interpretation. PCAs were performed separately for the PAL behavioral data and the histological data. For the behavior, only PAL measures were included, as no conclusive effects of injury condition were found for iPAL. For each animal, behavioral measures included the average percent correct, location bias, stimulus bias, and stimulus lose-shift index for the first five days of PAL as well as the number of errors made until criterion was met. Only the first component of this analysis had an eigenvalue above 1. This component explained over 88% of the variance in the dataset, with each of the five measures loading comparably between |0.38-0.48|. Of note, percent correct had a positive loading and each of the other four performance measures loaded negatively, exemplifying a prior known relationship. Component scores for each animal were then taken from this first and only significant component for subsequent correlation analysis.

For the histological data, measures for each animal included an average of the optical density across ROIs for the following significant ANOVAs (as shown in Figures 6 and 7): GFAP ipsilateral hippocampus, GFAP contralateral hippocampus, GFAP ipsilateral thalamus, IBA1 ipsilateral thalamus, & IBA1 contralateral thalamus. Spared tissue volume measurements for the cortex and hippocampus were also included to encompass the spread of data that contributed to the significant hemisphere volume analysis. This analysis produced three components with an eigenvalue above 1. The first principal component had high loadings for the GFAP staining and accounted for 42.05% of the total variance. The second principal component had high loadings for the IBA1 staining and accounted for 28.18% of the total variance. The third component corresponded to the spared tissue volume of the hippocampus and cortex. This component explained 16.20% of the total variance. Together, these three components explained over 82% of the variance in this data set. Table 1 shows the variable loadings on these three components. As with the behavioral PCA, the component scores for each animal across these three components were taken for subsequent correlation analysis. A graphical depiction of the component dimensions and clustering seen with this analysis is depicted in Figure 8.

**Table 1.**
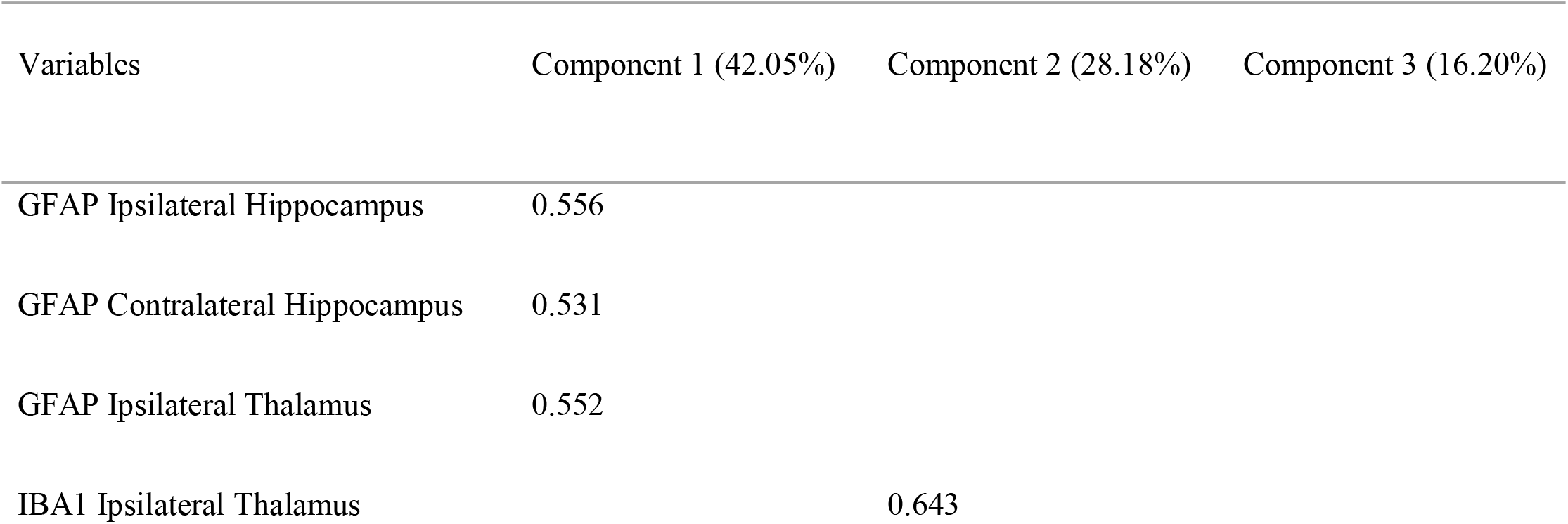
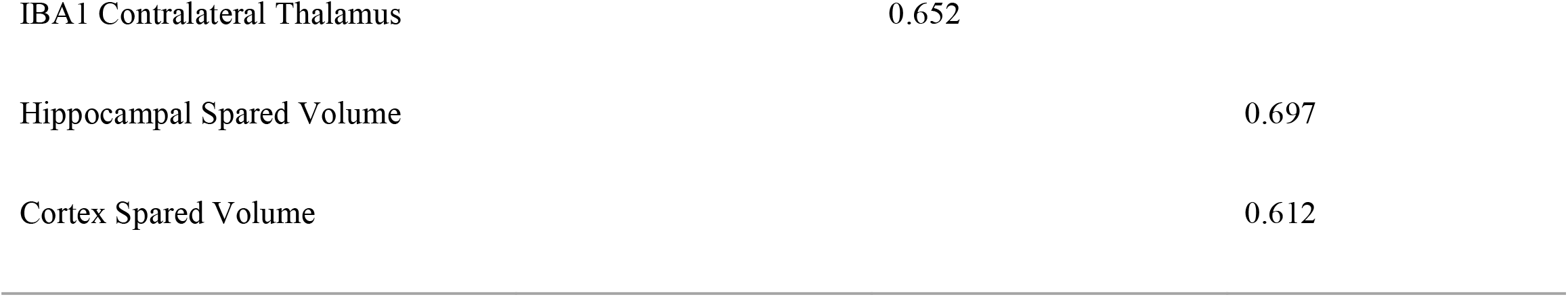
Loading values for the seven variables of the histology PCA analysis. Component 1 (42.05% of variance) corresponded to GFAP staining as indicated by the high loadings for the three GFAP variables, component 2 (28.18% of variance) corresponded to IBA1 staining, and component 3 (16.2% of variance) corresponded to spared tissue volume. Only loadings above 0.4 are depicted.

**Figure 8.**
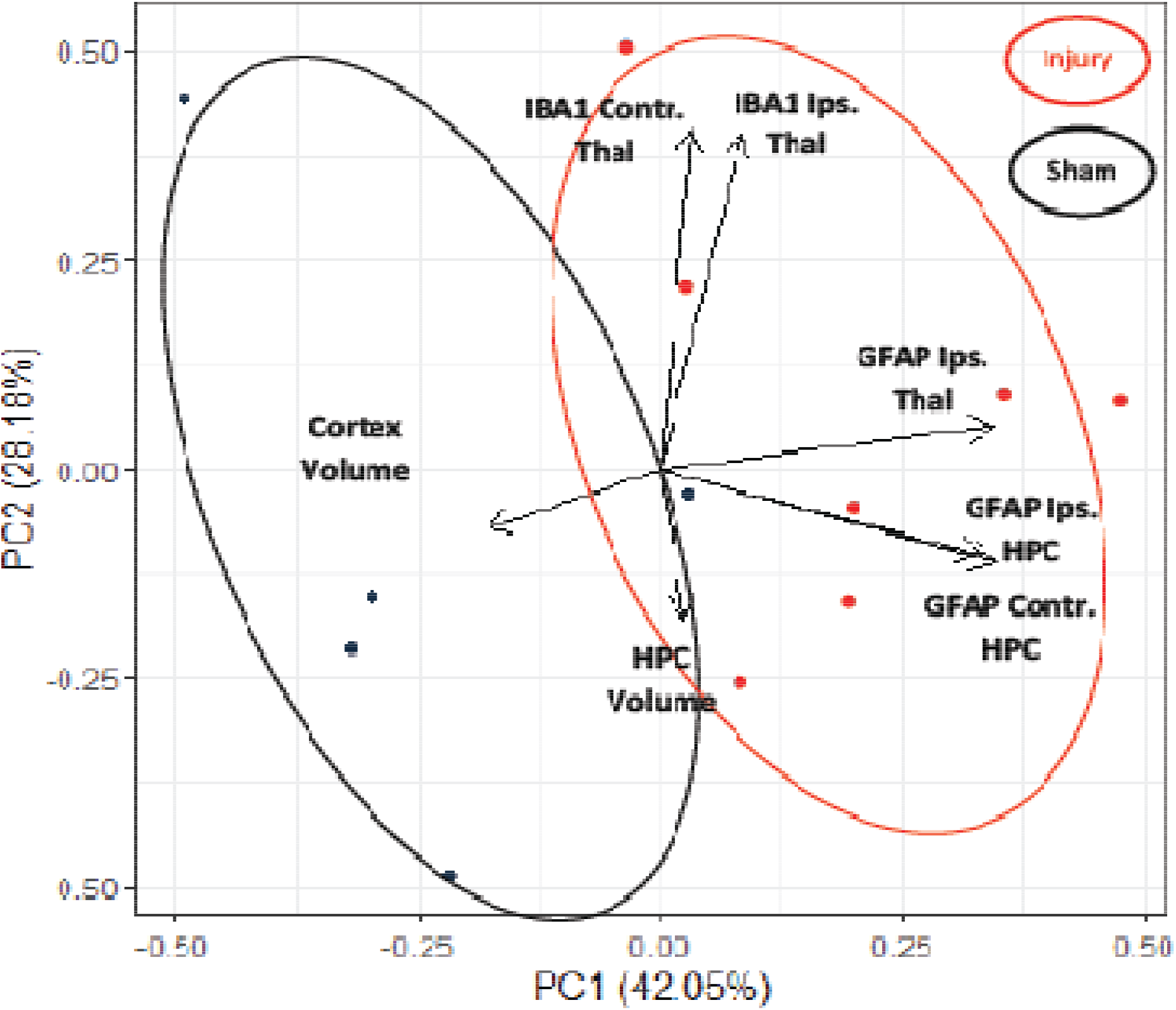
Graphical representation of the component dimensions and clustering for the histology PCA analysis. Although the model was naïve to injury condition, the histology PCA clustered CCI and sham animals into two distinct groups. Three components above an eigenvalue of 1 were found.

Taking the 4 component scores obtained for each animal (1 behavioral and 3 histological), we calculated Pearson’s correlation coefficients (Table 2). This analysis revealed a significant correlation between the PAL behavior and histology component 2, which corresponded to IBA1 staining in the thalamus (r = 0.83, p = 0.0008). Furthermore, there was also a significant correlation between histology component 3, which corresponded to spared tissue volume, and histology component 1, which corresponded to GFAP staining (r = 0.602, p = 0.039). When accounting for multiple comparisons, however, the adjusted p-value is nonsignificant. Finally, the IBA1 and GFAP components displayed a highly significant correlation (r = 0.990, p < 0.0001). This was expected, but it is particularly noteworthy these highly correlated components did not display the same correlation to the PAL behavior (Fisher z-transformation; z_(12)_ = 1.733, p = 0.08.)

**Table 2.**
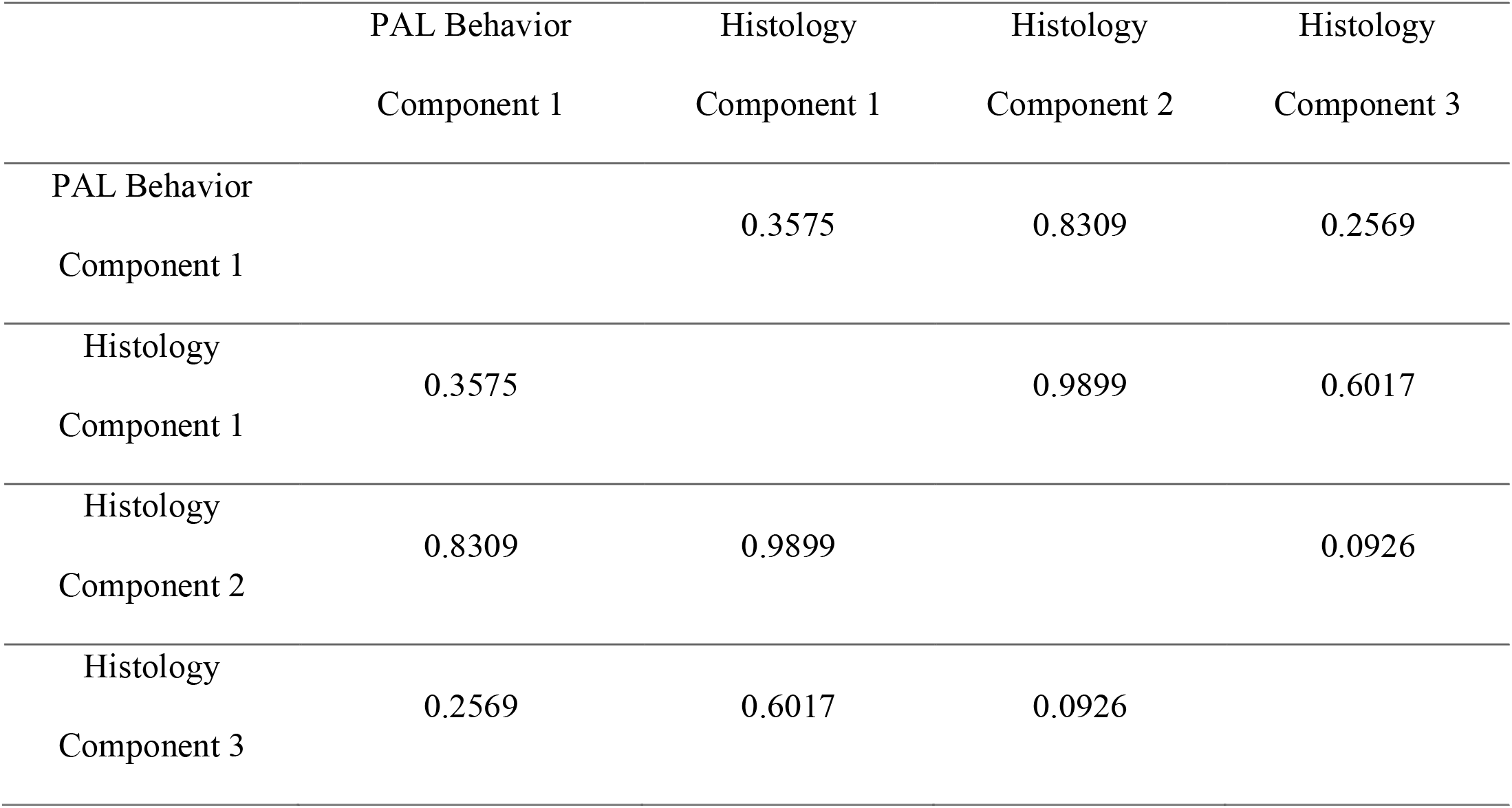
Pearson’s correlation coefficients for the four PCA components (one behavior and three histology). PAL behavior was highly correlated with IBA1 staining in the thalamus (p<0.001). Spared tissue volume was moderately corrrelated with GFAP staining, but was statistically nonsignifcant after correction for multiple comparisons (p>0.05). IBA1 and GFAP components were extremely correlated (p<0.0001), and it is noteworthy that these highly correlated components did not display the same correlation to PAL behavior (p=0.08).

## 4. DISCUSSION

The current study reports the deficits observed on a clinically relevant behavioral task, Paired Associates Learning (PAL), in a rat controlled cortical impact (CCI) model of traumatic brain injury (TBI). In addition to detailed behavioral performance measures, post-mortem histology was implemented to quantify GFAP/IBA1 immunolabeling and spared tissue volume. Principal component analysis was then used for dimension reduction to further understand the relations between the histological markers of TBI and the behavioral measurements obtained in PAL. Despite visible variability in the tissue loss following TBI and immunostaining (Figures 5,6,7), PAL performance was found to decline in the injury condition after CCI compared to the sham condition (Figure 3). Furthermore, after reducing several PAL performance measures into a single principal component and the histological measures into three principal components, there was a significant correlation of the rats’ component scores between the PAL behavior and the principal component related to thalamic IBA1 staining in the histology (Figure 8).

Our results validate previous findings from human study participants that PAL is sensitive to detecting cognitive impairment after TBI (Newcomb et al. 2011; Simoni et al. 2016). Furthermore, our detailed analysis of behavioral deficit on PAL post-TBI compliments a recent study in rats using PAL as a cognitive training paradigm after TBI (Braeckman et al. 2020). Braeckman et al. found that cognitive training on PAL increased microstructural organization in the hippocampus as measured by diffusion tensor imaging (DTI) metrics compared to no cognitive training. In our study, subcortical structures below the injured cortical region, namely a large portion of the hippocampus and midline thalamic nuclei, were shown to have accumulated protein markers of inflammatory processes. The hippocampus is integral to PAL performance in rats (Talpos et al. 2009; Langston et al. 2010; Yoon et al. 2012), suggesting that the observed impairments in PAL performance in the injured rats could be due to damage in this structure. Furthermore, altered resting state functional connectivity between the parietal cortex and hippocampus has been shown after TBI in rats (Mishra et al. 2014), and may be related to the deficits described here. As for the density of staining in the thalamic nuclei, there is extensive literature demonstrating that blood brain barrier dysfunction is prominent after TBI and may largely affect the thalamus (Ramlackhansingh et al. 2011; Vliet et al. 2020). Furthermore, circulating GFAP and IBA1 levels were recently found to be associated with pathophysiological sequelae in the thalamus of a pig model of mild TBI (Lafrenaye et al. 2020).

While our ROIs did not specifically include the anterior thalamic nucleus (ATN) due to the location of injury site, it is likely that blood brain barrier disruption and inflammatory glial response also affected the ATN. Furthermore, the ATN is densely reciprocally connected to the hippocampus and likely interacts with medial temporal structures to support associative learning (Savage, Hall, & Vetreno 2011; Perry et al. 2018; Sziklas & Petrides 1999). It is well established that the primary cause of secondary injury post-TBI is neuroinflammation (Giunta et al. 2012; Acosta et al. 2013; Simon et al. 2017). While controlling neuroinflammation appears to be a strong candidate for TBI therapeutics, some current anti-inflammatory treatments have demonstrated little to no cognitive benefits in clinical testing (Homsi et al. 2009; Kelso et al. 2011; Ding et al. 2014). Our findings that PAL task performance significantly correlated with IBA1 staining, a measure related to neuroinflammation, underscores PAL as a well-suited task for preclinical testing of anti-inflammatory interventions for TBI.

Our study is not without limitations and interpretation of our results should be made within the appropriate context. First, the controlled cortical impact model of TBI used in this study likely doesn’t reflect the average injury seen in the human population. Future studies may benefit from the use of more relevant injury models such as that of the closed-head model of engineered rotational acceleration (Harr et al. 2019; McNamara et al. 2020). While we did assess the extent of lesion volume through the analysis of spared tissue, other studies have shown that structural white matter damage after TBI can reflect neurological outcomes (Kinnunen et al. 2011; Newcombe et al. 2016; Braun et al. 2017). Furthermore, Braeckman et al. (2020) showed that DTI metrics were a reliable measure to evaluate cognitive training in their rat model of TBI. Employing DTI with the trial-by-trial analysis of PAL deficits in this study will be a promising avenue. Moreover, our study was not designed to have consistent endpoints for all rats. Some rats took longer to complete PAL than others, and thus the time from post-surgery to sacrifice differed slightly between rats. Future studies using this model will benefit from a time-restricted analysis of microglia and astrocyte density after TBI. Finally, we were not sufficiently powered to detect sex differences, though we reported the addition of sex to our LMEMs.

## 5. CONCLUSIONS

Understanding the complexities of TBI requires an evolving and reciprocal relationship between basic, translational, and clinical biomedical research. The development of automated touchscreen testing not only affords researchers a range of detailed behavioral data to assess performance more accurately on cognitive tasks, but also gives translational research the power to test methods of intervention in clinically relevant settings. This study found PAL task performance to decline after a unilateral parietal lobe impact that resulted in greater than expected injury variation, emphasizing PAL as a useful tool for testing the efficacy of interventions in preclinical models of TBI. The successful treatment of TBI will likely rely upon individualized therapies that take into consideration the variety of factors that can influence positive treatment outcomes and the thorough behavioral analyses presented here for the PAL task may be useful for unpacking individual differences during intervention testing for TBI. We believe this study closes a research gap with a level of transparency and breadth that will be important for future research.

## Acknowledgements

This research was funded by the McKnight Brain Research Foundation Institute, the Department of Defense GRANT11811993 for the impact of PERK on post-traumatic tauopathy in Alzheimer’s disease (JA), NIA R01AG049722/2RF1AG049722 (SNB), and the University of Florida Summer Neuroscience Internship Program (SMS). The authors would like to thank Aleyna Ross for the PAL graphic, Sabrina Zequeira for her help in the development of the iPAL task, and Wonn Pyon for helpful coding and analytical discussion.

