## Supplementary material for "Paired Associates Learning is Disrupted After Unilateral Parietal Lobe Controlled Cortical Impact in Rats: A Trial-by-Trial Behavioral Analysis": Figures

### Slide 1
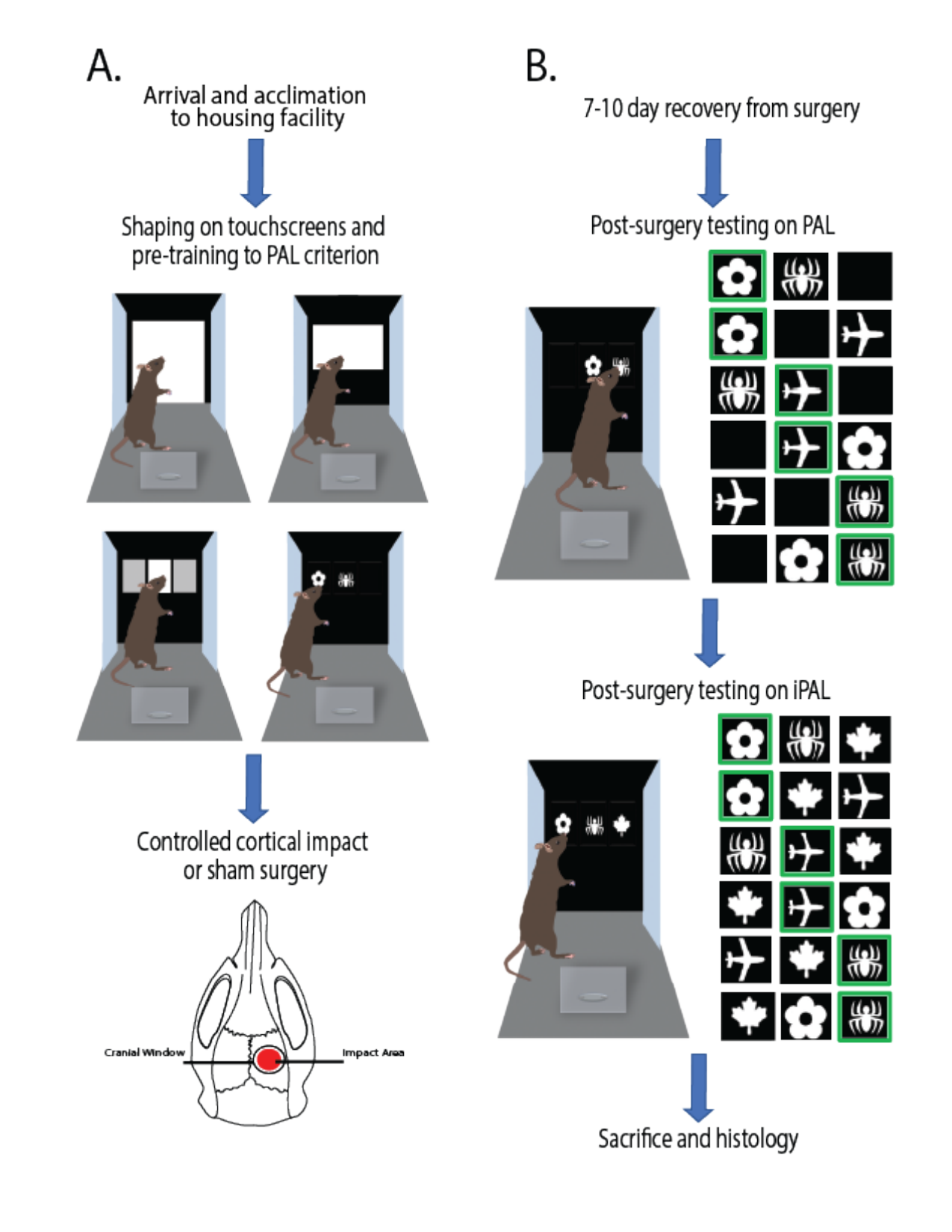

### Slide 2
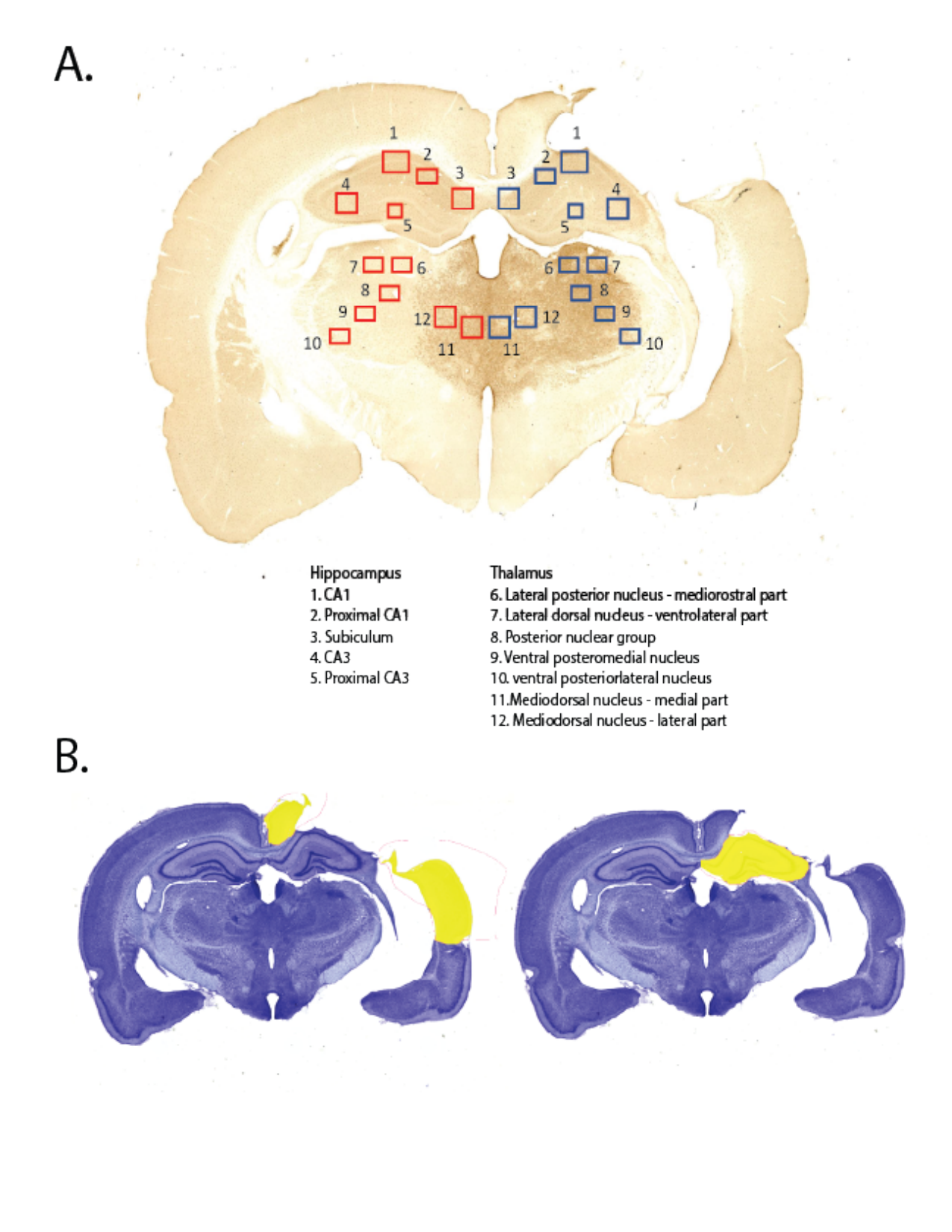

### Slide 3
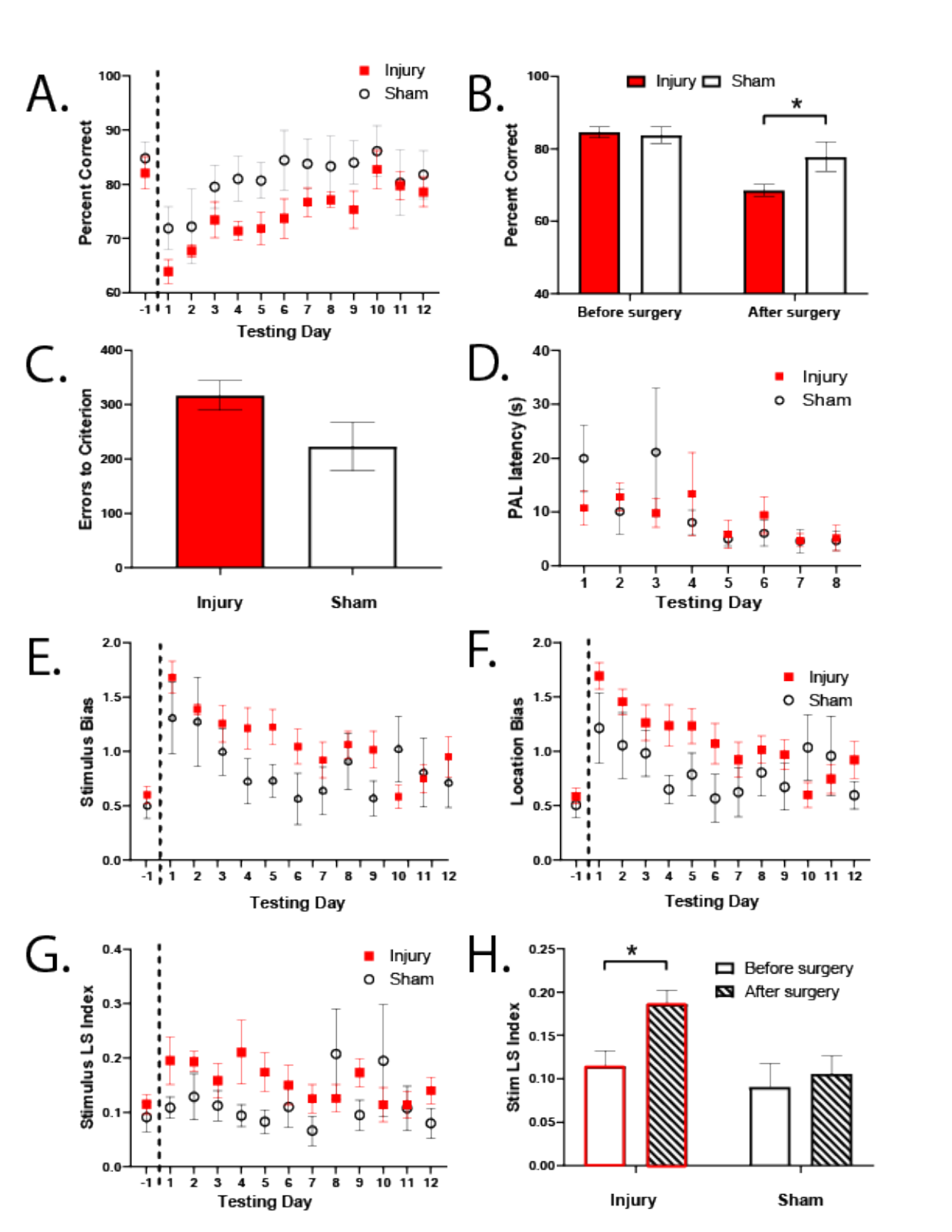

### Slide 4
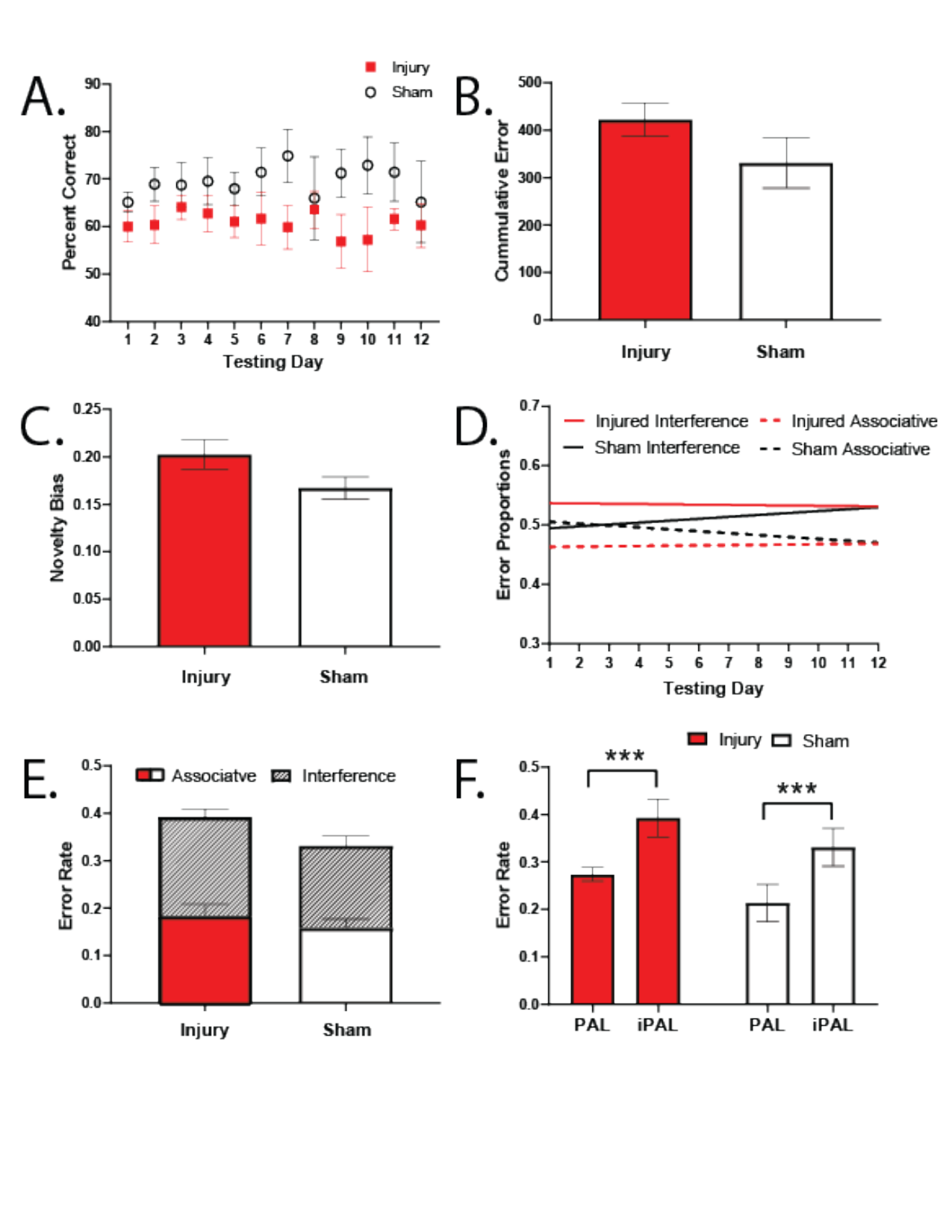

### Slide 5
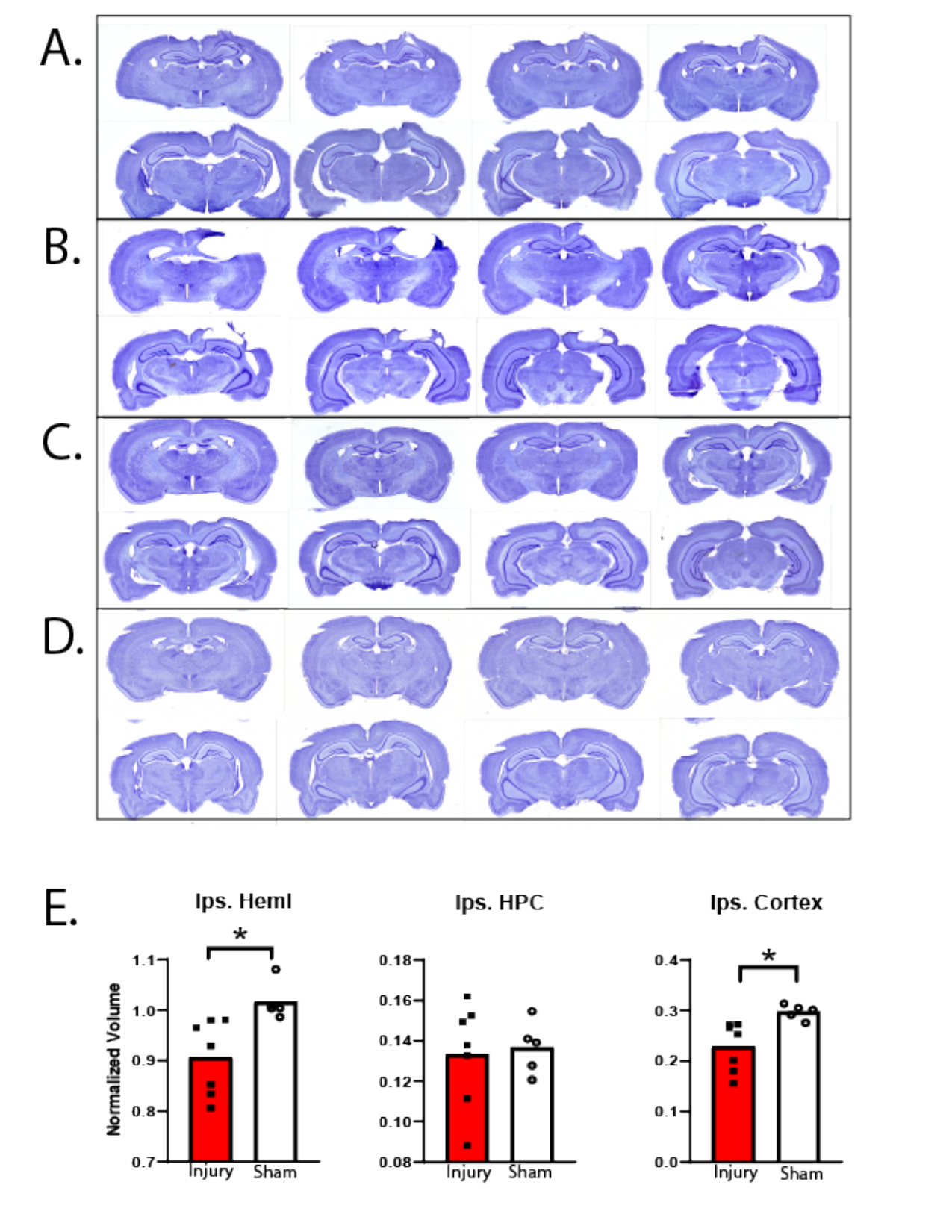

### Slide 6
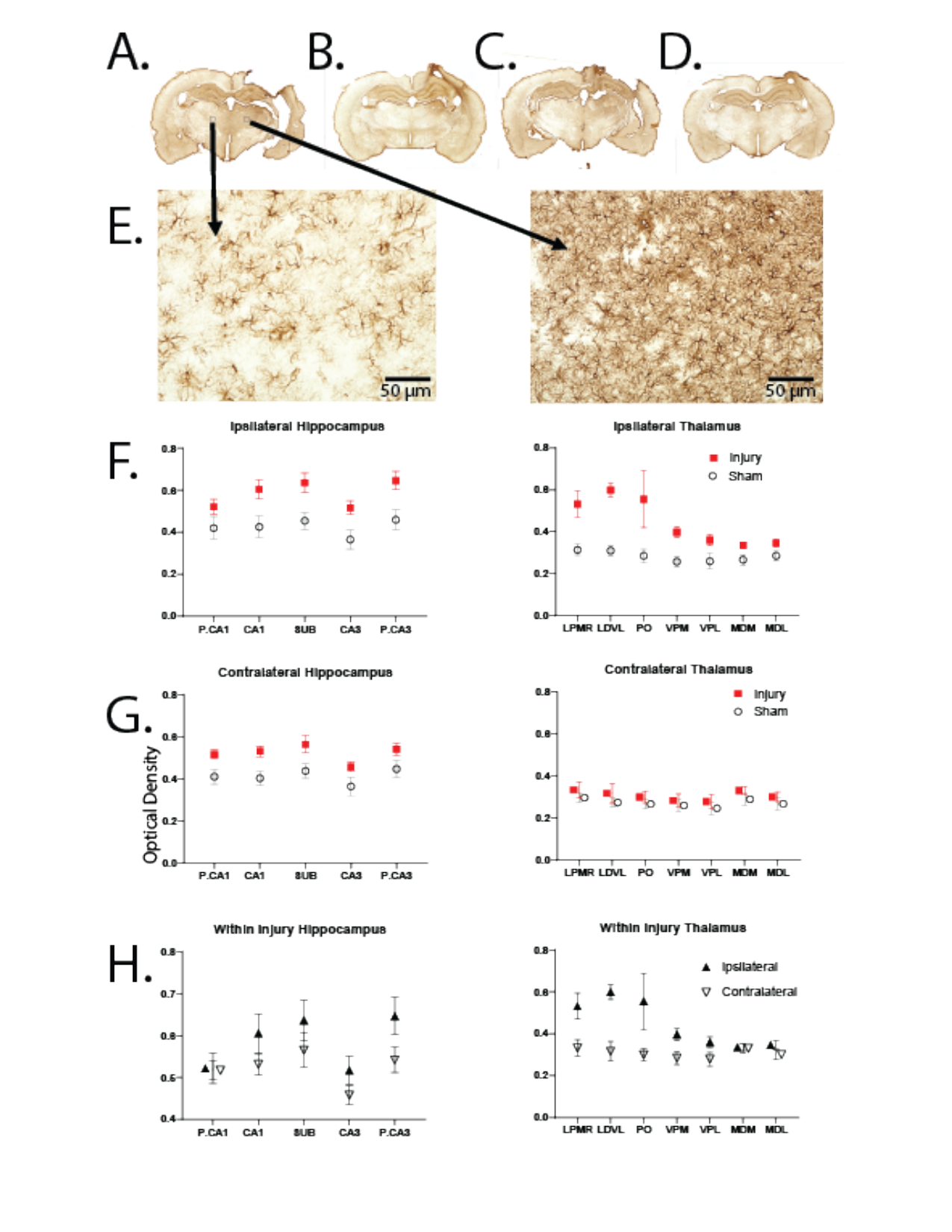

### Slide 7
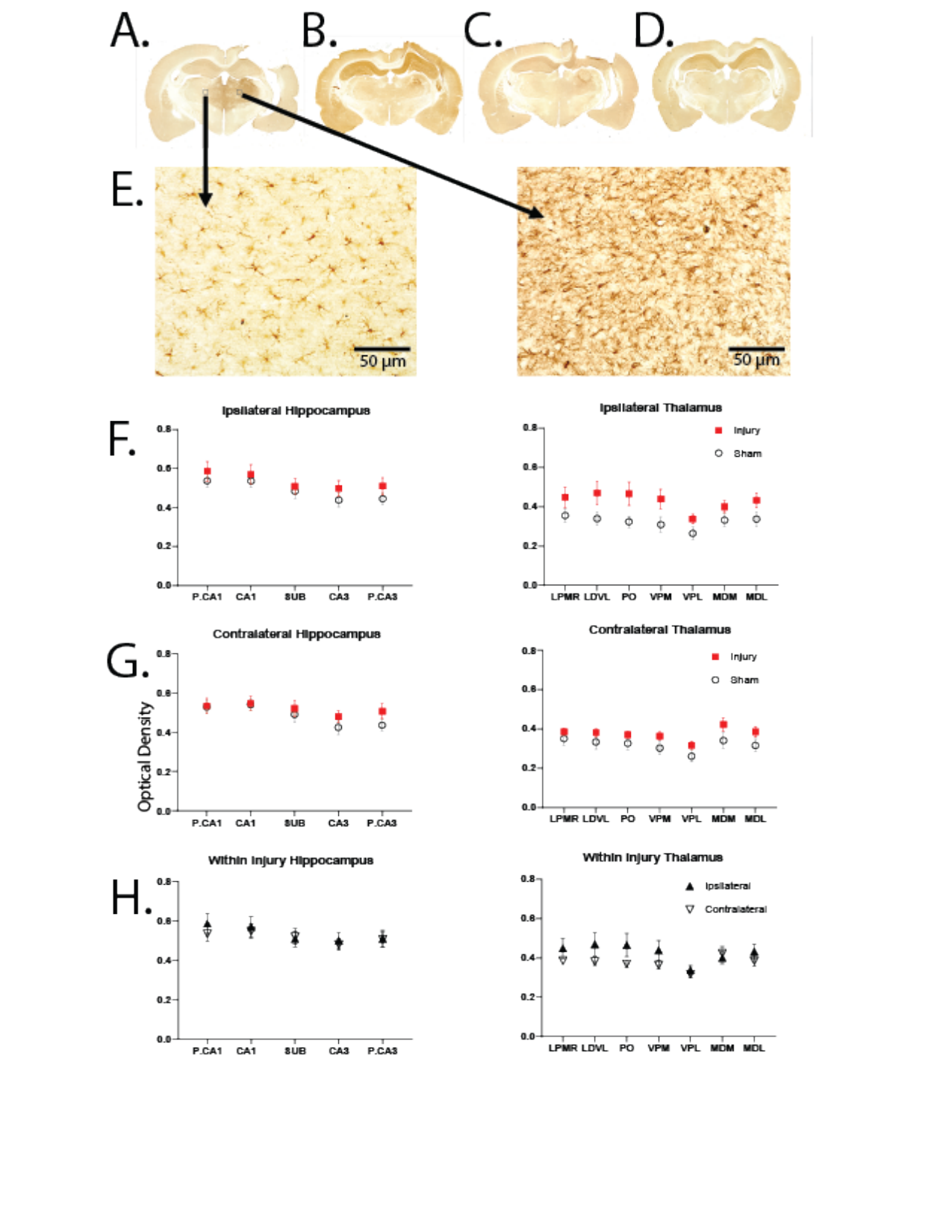

### Slide 8
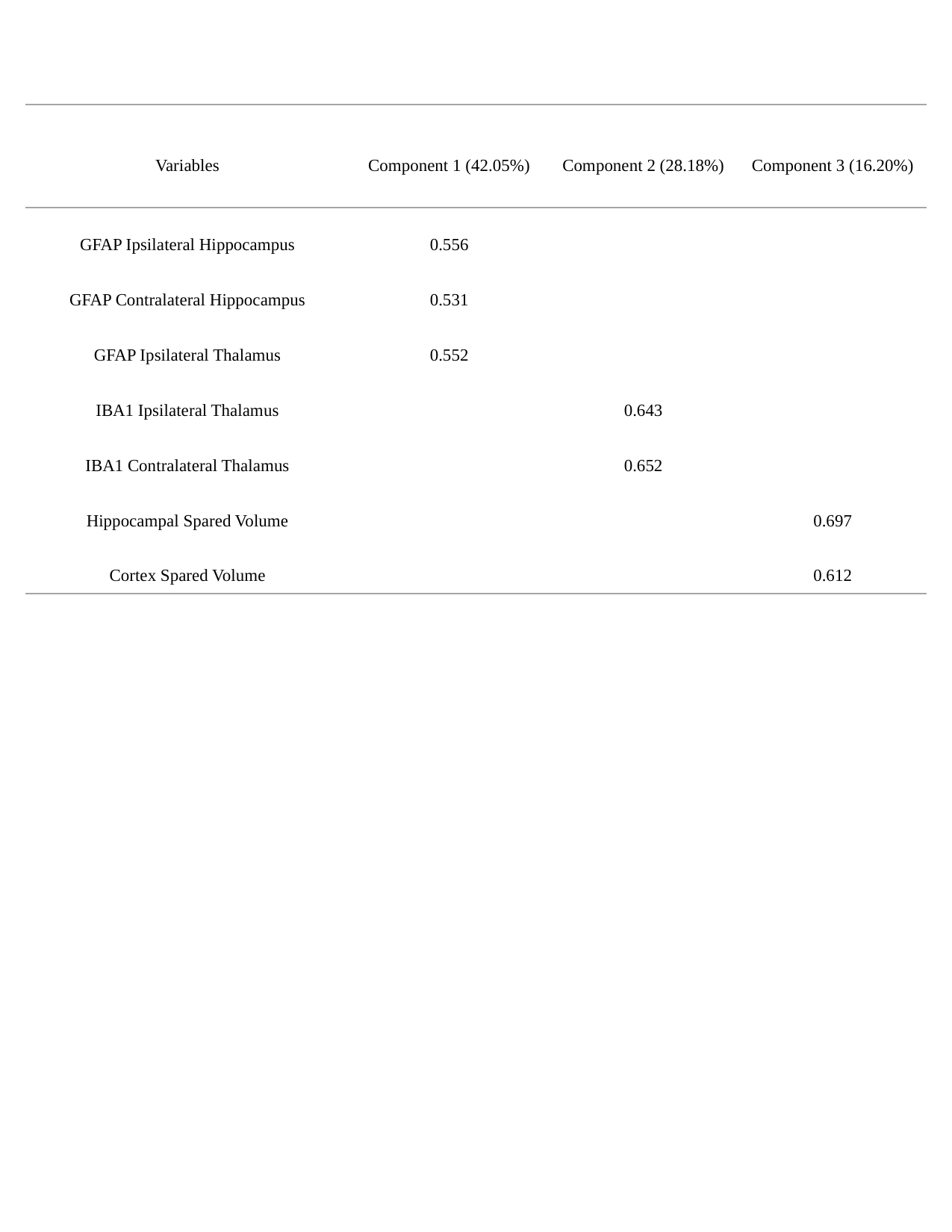

| Variables | Component 1 (42.05%) | Component 2 (28.18%) | Component 3 (16.20%) |
| --- | --- | --- | --- |
| GFAP Ipsilateral Hippocampus | 0.556 | | |
| GFAP Contralateral Hippocampus | 0.531 | | |
| GFAP Ipsilateral Thalamus | 0.552 | | |
| IBA1 Ipsilateral Thalamus | | 0.643 | |
| IBA1 Contralateral Thalamus | | 0.652 | |
| Hippocampal Spared Volume | | | 0.697 |
| Cortex Spared Volume | | | 0.612 |

### Slide 9
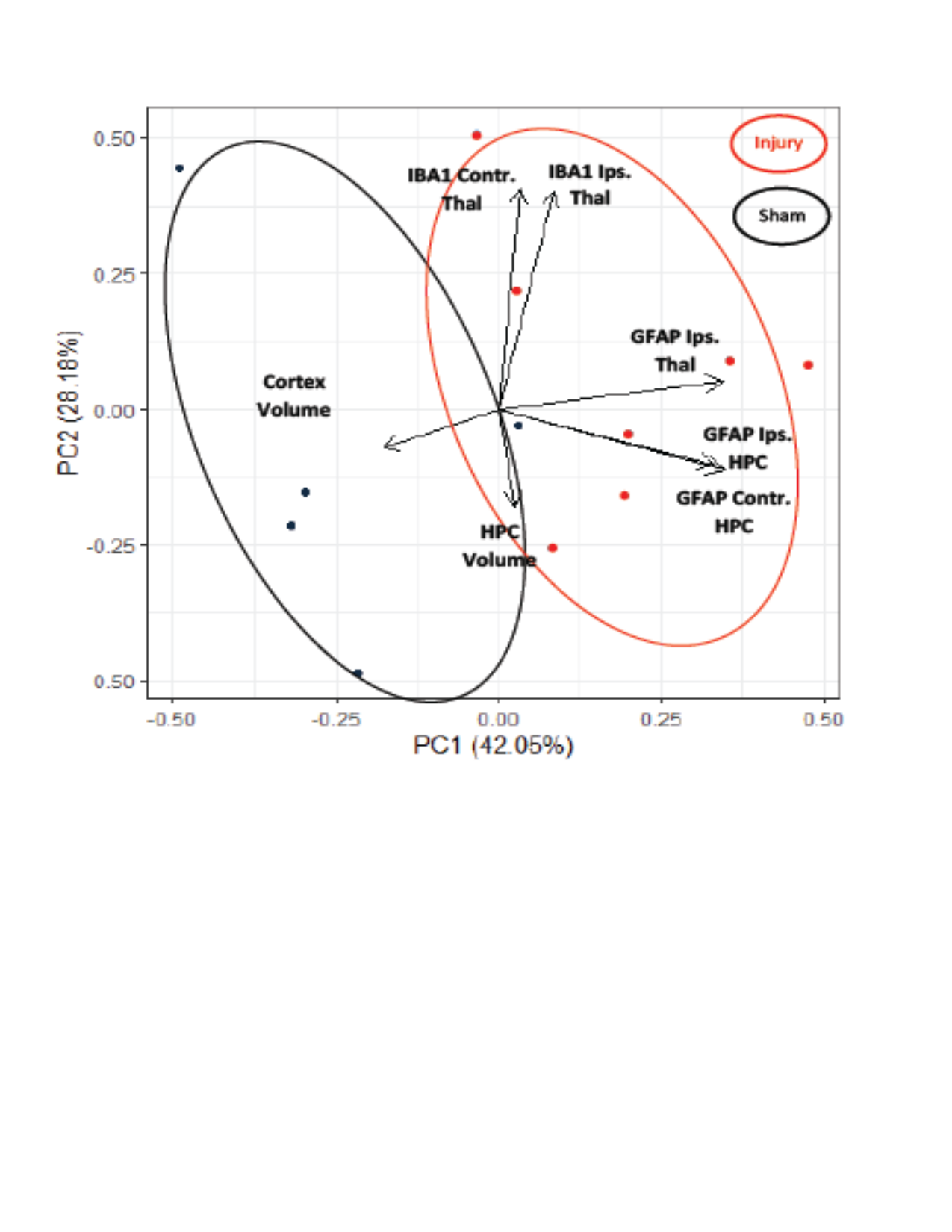

### Slide 10
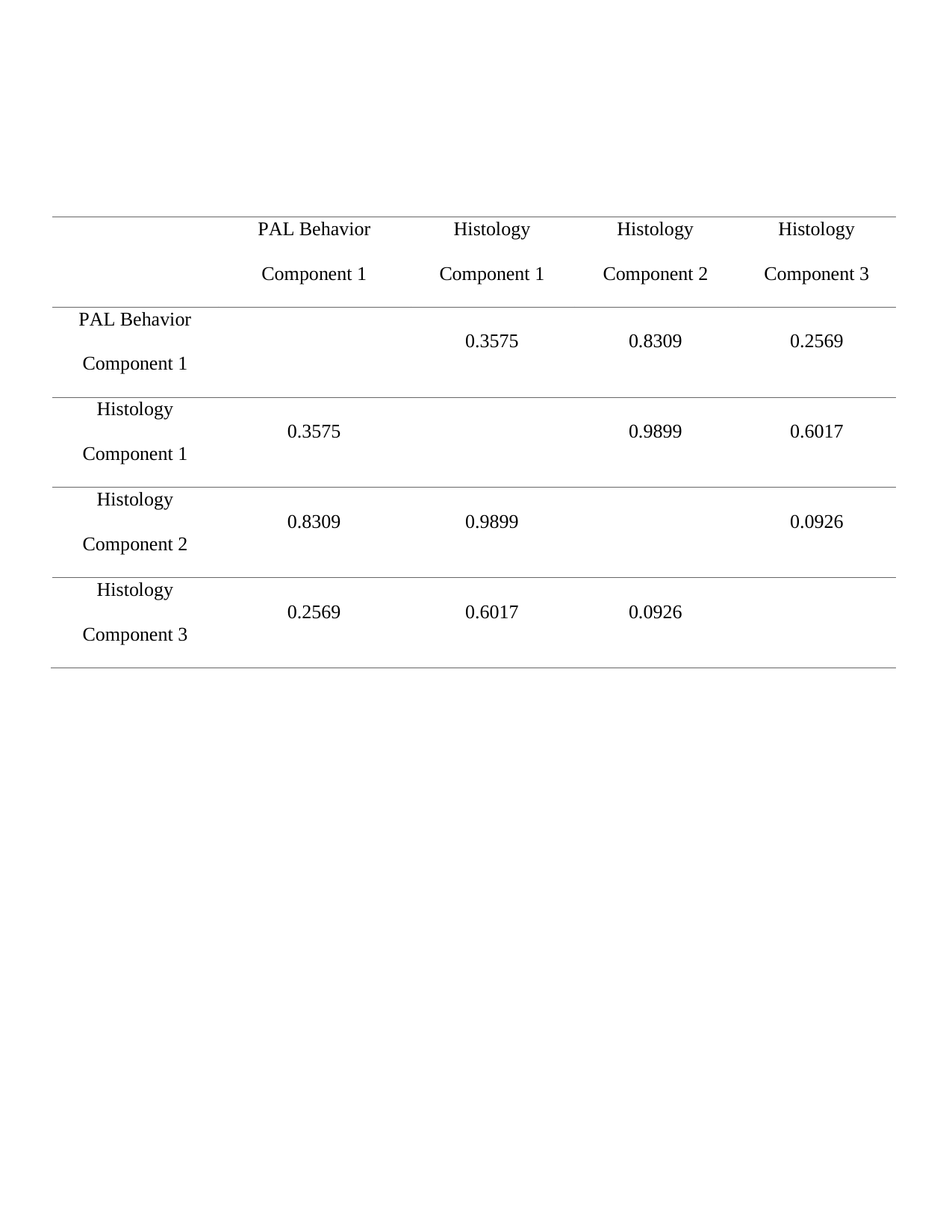
